# *In situ* X-ray assisted electron microscopy staining for large biological samples

**DOI:** 10.1101/2021.06.19.448808

**Authors:** Sebastian Ströh, Eric W. Hammerschmith, David W. Tank, H. Sebastian Seung, Adrian A. Wanner

## Abstract

Electron microscopy of biological tissue has recently seen an unprecedented increase in imaging throughput moving the ultrastructural analysis of large tissue blocks such as whole brains into the realm of the feasible. However, homogeneous, high quality electron microscopy staining of large biological samples is still a major challenge. To date, assessing the staining quality in electron microscopy requires running a sample through the entire staining protocol end-to-end, which can take weeks or even months for large samples, rendering protocol optimization for such samples to be inefficient.

Here we present an *in situ* time-lapsed X-ray assisted staining procedure that opens the “black box” of electron microscopy staining and allows observation of individual staining steps in real time. Using this novel method we measured the accumulation of heavy metals in large tissue samples immersed in different staining solutions. We show that the measured accumulation of osmium in fixed tissue obeys empirically a quadratic dependence between the incubation time and sample size. We found that potassium ferrocyanide, a classic reducing agent for osmium tetroxide, clears the tissue after osmium staining and that the tissue expands in osmium tetroxide solution, but shrinks in reduced osmium solution.

X-ray assisted staining gave access to the *in situ* staining kinetics and allowed us to develop a diffusion-reaction-advection model that accurately simulates the measured accumulation of osmium in tissue. These are first steps towards *in silico* staining experiments and simulation-guided optimization of staining protocols for large samples. Hence, X-ray assisted staining will be a useful tool for the development of reliable staining procedures for large samples such as entire brains of mice, monkeys or humans.

## Introduction

In the past decade the image acquisition rates of biological electron microscopy facilities have been scaled to 10^7^ – 10^9^ pixels per second through parallelization and automation(Hayworth et al. 2014; Schalek et al. 2011; Ren and Kruit 2016; Denk and Horstmann 2004; Hayworth et al. 2020; Eberle et al. 2015; Graham et al. 2019; Xu et al. 2017). The increase in imaging throughput for biological samples has been mainly driven by the emerging Neuroscience field of connectomics which aims to densely reconstruct neuronal circuits with synaptic resolution(Briggman, Helmstaedter, and Denk 2011; Helmstaedter et al. 2013; Bock et al. 2011; Zheng et al. 2018; Scheffer et al. 2020; Kornfeld et al. 2017; Wanner and Friedrich 2020; Lee et al. 2016; Kasthuri et al. 2015; Wilson et al. 2019; Schmidt et al. 2017; Svara et al. 2018; Vishwanathan et al. 2017). Also, the analysis of the terabyte-sized electron microscopy datasets produced by these studies is becoming increasingly automated using machine-learning and computer vision(Januszewski et al. 2018; Dorkenwald et al. 2020; Schubert et al. 2019; Dorkenwald et al. 2017; Berning, Boergens, and Helmstaedter 2015; Staffler et al. 2017; Jain, Seung, and Turaga 2010; Buhmann et al. 2020; Vergara et al. 2020; Turner et al. 2020). Both the image acquisition as well as the image analysis for these types of datasets are being scaled by parallelization to large samples on the order of several cubic millimeters or even entire brains. However, a remaining obstacle is the lack of reliable tissue processing and staining protocols for large samples where the smallest dimension is greater than 1 mm.

Existing *en bloc* electron microscopy staining protocols have been optimized for staining small samples with dimensions of less than 1 mm(Genoud et al. 2018; Hua, Laserstein, and Helmstaedter 2015; Deerinck et al. 2010; Tapia et al. 2012). Using aldehyde-stabilized cryopreservation(McIntyre and Fahy 2015) the cellular ultrastructure can be preserved even in large tissue blocks with dimensions exceeding 1 mm and entire brains. Despite pioneering work on *en bloc* staining protocols for whole mouse brains(Mikula and Denk 2015; Mikula, Binding, and Denk 2012), large *en bloc* stained samples still suffer from artifacts such as inhomogeneous staining and membrane or tissue cracks.

*En bloc* sample preparation for electron microscopy generally requires tissue fixation, staining and embedding in resin. First, the macromolecules in the tissue are stabilized via crosslinking by diffusing or perfusing buffered solutions of fixatives such as paraformaldehyde and glutaraldehyde (Claude and Fullam 1945; Fox et al. 1985; Sabatini, Miller, and Barrnett 1964). Because biological tissue is composed mostly of carbon and other low atomic number elements, the tissue is stained with heavy metals to increase the contrast in electron micrographs(Bahr 1954; Porter, Claude, and Fullam 1945). These heavy metal stains are composed of electron dense atoms such as osmium, lead or uranium. In addition, some of these heavy metals (e.g. osmium) also act as fixatives (Bahr 1955). Finally, the stained tissue gets dehydrated and embedded in resin. The most commonly used resins are epoxy resins(Maaløe O And Birch-Andersen 1956), such as araldite(Glauert, Rogers, and Glauert 1956) or Epon(Finck 1960) because of their thermal stability and electron transparency. To inspect the staining quality in an electron microscope, ultrathin sections (<100 nm) are typically collected from the embedded tissue using an ultramicrotome.

Many steps in the classic *en bloc* electron microscopy staining protocols are based on passive diffusion of chemicals into the tissue. Passive diffusion is one of the main bottlenecks for preparing large samples (smallest dimension >1mm) for electron microscopy(Burkl and Schiechl 1968; Medawar 1941). For small samples (smallest dimension <1mm) a typical *en bloc* staining protocol takes about 10 – 15 days including sectioning(Tapia et al. 2012). However, for large samples or whole brains, a diffusion-based staining protocol takes several weeks or even months(Mikula, Binding, and Denk 2012; Mikula and Denk 2015; Masís et al. 2018).

To date, electron microscopy staining is a “black box” and changes to staining protocols can only be assessed using an electron microscope, which in turn requires a sample to be run through the entire staining protocol. Conventional approaches for optimizing the parameters of staining protocols rely on sequential screening of hundreds of samples that have been processed end-to-end. However, sequential screening is very inefficient for months-long protocols.

Recently, X-ray based computed micro-tomography (μCT) has been introduced for a relatively fast, macroscopic assessment of staining quality and tissue integrity of resin-embedded whole mouse brains(Mikula and Denk 2015; Kuan et al. 2020; Dyer et al. 2017). Building on this pioneering work, we developed *in situ* time-lapsed X-ray assisted staining to observe the staining process while the samples are in the staining solution (Figure 1a). We used X-ray assisted staining to explore the micro-scale tissue mechanics and the kinetics of the heavy metal diffusion and accumulation in large aldehyde-fixed brain tissue blocks, resulting in new insights on how different staining agents affect the tissue. X-ray assisted staining opens the “black box” of electron microscopy staining protocols. Each staining step can be monitored and assessed in real time. This enables *in silico* optimization of electron microscopy staining protocols, which will be particularly useful for the development of staining procedures for large biological samples such as whole brains (Figure 1g).

**Figure 1:**
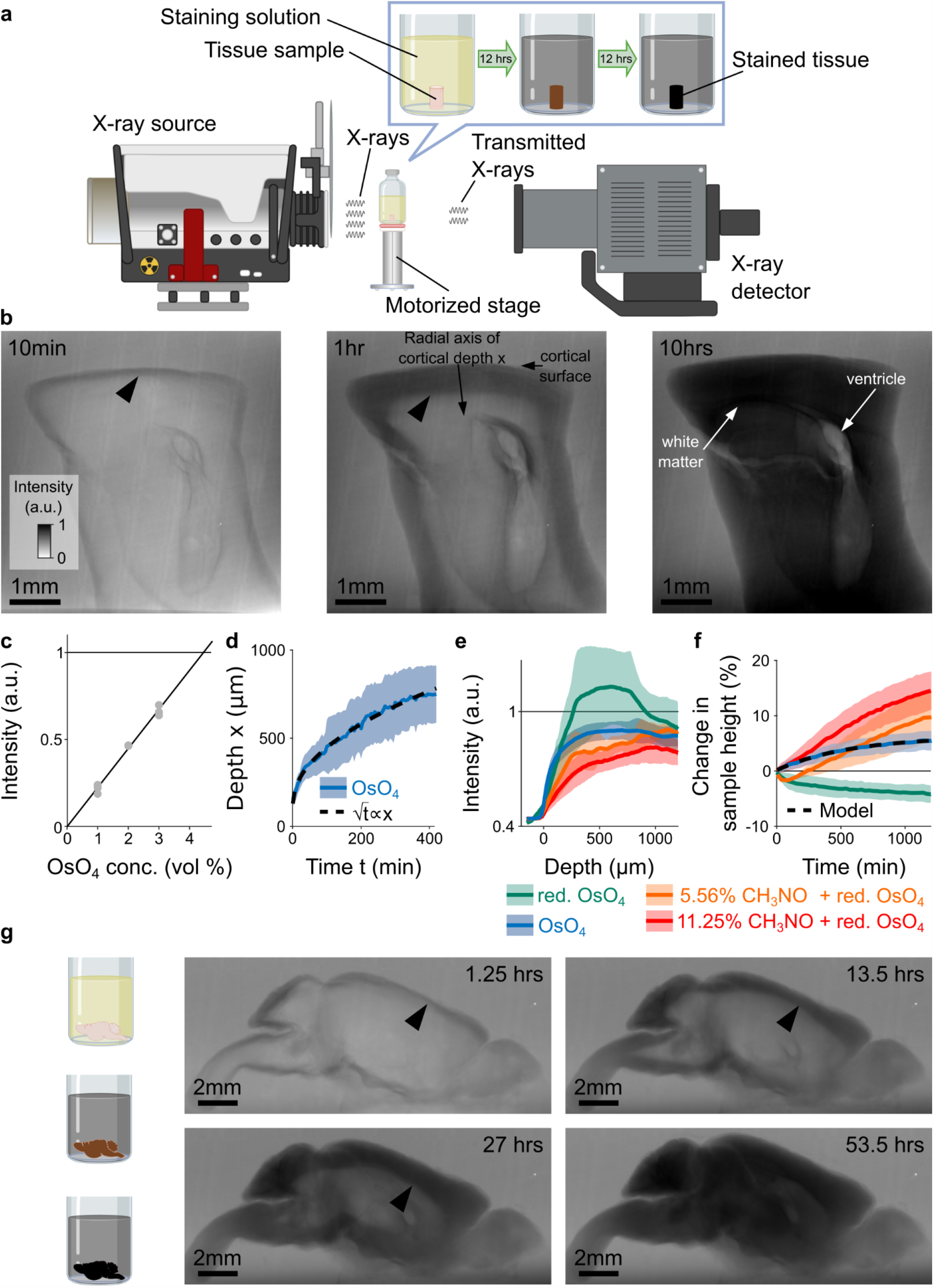
X-ray assisted staining. **a** Experimental setup. The sample sits in staining solution in a sealed glass vial on a motorized stage. X-rays are emitted from an X-ray source, pass through the sample, and the spatial distribution of the transmitted X-rays is detected and used to form a projection image of the X-ray absorption in the sample. Over time, the heavy metals diffuse into the tissue and the accumulation of heavy metals change the X-ray absorption properties of the sample. For the quantitative results presented in this study we used cylindrical tissue samples extracted with a 4-mm diameter biopsy punch from the cortex of mice. **b** Projection images of a 4-mm punch of mouse cortex incubated in 2% buffered osmium tetroxide (OsO_4_) solution after 10 min, 1 hr and 10 hrs. The osmium diffuses passively from the cortical surface and the brain ventricles into the tissue and forms a staining front of tissue-bound osmium (arrowhead) that moves towards the center of the sample as time progresses. The accumulation of heavy metals results in an increase in the pixel intensity (a.u.) of the X-ray projection image, which was measured along the radial axis of cortical depth from the cortical surface towards the white matter. **c** Measured pixel intensities of the X-ray projection images for different concentrations of buffered OsO_4_ solutions. The higher/darker the intensity, the more X-rays are absorbed. The pixel intensity/X-ray absorption scales linearly with the concentration/density of osmium (black line, Pearson correlation coefficient r=0.99, p<10^−6^; n=3 for each concentration). **d** Propagation of the staining front in 4-mm brain punches immersed in 2% buffered OsO_4_ solution (n=13) measured along the radial axis of cortical depth (mean ± s.d.). The propagation can be fitted by a quadratic model (dashed black lines) in which the penetration depth x of the staining front is proportional to the square root of the incubation time t (residual standard error: SE_res_=14.68 μm). **e** Intensity profiles of the spatial heavy metal accumulation after t=20 hours of incubation in 2% buffered OsO_4_ (blue, n=13), reduced OsO_4_ (green, n=6). reduced osmium with 5.56% (orange, n=4) and 11.25% formamide (red, n=4) solutions (mean ± s.d.). In reduced osmium the heavy metal staining is not homogenous and there is a more densely stained tissue band at a depth of 300 – 800 μm. **f** Tissue expansion quantified as change in sample height of aldehyde-fixed brain punches immersed in OsO_4_ (blue, n=13), reduced OsO_4_ (green, n=6). reduced OsO_4_ with 5.56% (orange, n=4) and 11.25% formamide (red, n=4), respectively, as a function of the immersion duration (mean ± s.d.). All solutions were buffered by cacodylate. A monomolecular growth model was fitted to the average OsO_4_ expansion (black dashed line, residual standard error SE_res_=0.06%). **g** *In situ* X-ray assisted staining of large tissue samples. The images show projections of a whole mouse brain incubated in 2% buffered OsO_4_ at different points in time across 2.5 days. The arrowheads indicate the location of the staining front of accumulated osmium. Scale bar 2 mm.

## Results

We developed *in situ* time-lapsed X-ray assisted staining with the goal of facilitating and accelerating the optimization of staining protocols for large biological samples such as whole mouse brains(Abbott et al. 2020). 4mm-brain punches of transcardially perfused mice were immersed in aldehyde fixatives for 36 hours. After washing the sample blocks with cacodylate buffer, the samples were immersed in staining solution and immediately placed in the acquisition chamber of a Zeiss Xradia 520 Versa 3D for X-ray microscopy (Figure 1a). The Xradia can be operated in two different modes:

- In the μCT mode, projections of the sample are acquired at different rotation angles in order to reconstruct a 3D computed tomograph of the sample. Depending on the required signal to noise and resolution, this mode allowed us to acquire approximately 1 – 2 computed tomograph per hour.
- In single-projection mode, a projection of the sample is acquired in a fixed position every few seconds without rotating the sample.

The advantage of the μCT mode is that arbitrary virtual reslices can be extracted to assess the staining progression in various parts of the sample. However, with our X-ray microscope, the temporal resolution in μCT mode was limited to 30 – 60 min per tomograph. Therefore we performed all quantitative experiments in single-projection mode where the temporal resolution for monitoring of the heavy metal diffusion and accumulation was on the order of a few seconds. As soon as the samples got placed in the staining solutions, the heavy metals started accumulating in the immersed tissue. The stained tissue absorbs more X-rays than the unstained tissue, resulting in a noticeable intensity difference that can be used to track the diffusion and accumulation of the heavy metals in the tissue (Figure 1b).

### Quadratic scaling of incubation times with sample size

Osmium tetroxide (OsO_4_) is one of the most commonly used contrast agents for lipid staining in electron microscopy due to its large atomic number and its ability to integrate into cellular membranes(Porter, Claude, and Fullam 1945; Palade 1952; Watson 1958). The immersion times for OsO_4_ vary between a few minutes to several days, depending on the sample size and tissue type, and are usually determined empirically(Genoud et al. 2018; Hua, Laserstein, and Helmstaedter 2015; Tapia et al. 2012; Mikula and Denk 2015). However, the kinetics of OsO_4_ staining of biological tissue are not well understood and so far had to be determined experimentally by trial and error. We therefore set out to measure the diffusion and accumulation of OsO_4_ by placing 4-mm punches of aldehyde-fixed mouse brains in 2% buffered OsO_4_ solution. The osmium diffused into the tissue and accumulated in the sample during the staining process (Figure 1b). The intensity of X-ray absorption in the projection images scales linearly with the local concentration/density of osmium (Figure 1c). The density of OsO_4_ in the tissue increased beyond the density in the surrounding staining solution, indicating that the density of binding sites for OsO_4_ in the tissue is higher than the concentration of the OsO_4_ in the staining solution (Figure 1b). The diffusion and spatio-temporal accumulation of OsO_4_ results in a staining front that propagates towards the center of the sample (Figure 1b). As expected for diffusive processes(Crank 1979), the propagation of the OsO_4_ staining front can be approximated by a quadratic model (Figure 1d). As in the case of other fixatives(Medawar 1941), the osmium staining penetration in aldehyde fixed tissue obeys a quadratic scaling law, or rather a “rule of thumb”, for how the necessary incubation time t depends on the staining depth or sample size x:

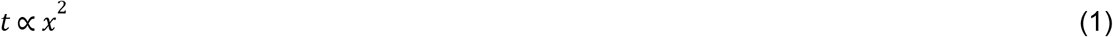

This means, for example, that if one would adapt an established OsO_4_ staining protocol for 3X larger samples, one would have to prolong the incubation time by 9X in order to produce comparable staining results. Similarly, the time it takes for any point in the sample to reach a given concentration is proportional to the square of its distance to the sample surface (Crank 1979).

### Monitoring staining kinetics and tissue deformation

After 20 hours of incubation the staining density of osmium is homogeneous across the first 1,000 μm of cortical depth (Figure 1b+e, Supplementary Figure 1). Reduced osmium is another commonly used staining agent that is known to result in higher contrast electron microscopy images than non-reduced OsO_4_. Typically, a buffered 2% OsO_4_ solution is reduced with 2.5% potassium ferrocyanide (K_4_[Fe(CN)_6_])(Hua, Laserstein, and Helmstaedter 2015; Willingham and Rutherford 1984; Mikula and Denk 2015). Consistent with previous reports on reduced osmium staining in large samples(Hua, Laserstein, and Helmstaedter 2015; Mikula and Denk 2015) we found that for reduced osmium the accumulation of heavy metals peaks at a depth between 300 – 800 μm (Figure 1e, Supplementary Figure 2), whereas the tissue above or below that depth is stained less. Traditionally, this band of more heavily stained tissue has been associated with precipitated osmium that hinders the diffusion and prevents homogenous staining(Genoud et al. 2018; Hua, Laserstein, and Helmstaedter 2015; Mikula and Denk 2015), in particular deeper in the tissue. Ref. (Mikula and Denk 2015) reported that homogeneous staining of large samples with reduced osmium was achieved by adding formamide to the reduced osmium solution. However, the underlying mechanisms by which formamide acts are not known(Mikula and Denk 2015; Genoud et al. 2018). Ref. (Mikula and Denk 2015) hypothesized that formamide might prevent precipitation by generally solubilizing compounds or that it might allow highly charged molecules to cross membranes more easily. Adding formamide to the reduced osmium solution indeed resulted in more homogeneous staining (Figure 1e, Supplementary Figures 3+4), but the heavy metal density was lower than in osmium only. Ref. (Mikula and Denk 2015) reported that the tissue tends to expand at high concentrations of formamide (>50%). But we found that the tissue expands even at lower formamide concentrations, while the amount of expansion depends on both the formamide concentration and the incubation duration (Figure 1f). For a concentration of 11.25% formamide, the sample height increased by about 15% within 20 hours of incubation, whereas for a formamide concentration of 5.56%, the tissue height expanded only by approximately 10% (Figure 1f). Note, in the first 3 hours of incubation in reduced osmium with 5.56% formamide the tissue actually shrank, suggesting that there are opposite forces acting on the tissue. Indeed, in reduced osmium solution without any formamide, the tissue shrank by about 5% in height (Figure 1f). In contrast, we found that in 2% buffered OsO_4_ solution the sample height expands by about 5% (Figure 1f).

Ref. (Hua, Laserstein, and Helmstaedter 2015) suggested an alternative approach to achieve homogeneous staining in 1-mm brain punches with reduced osmium without formamide, in which osmium and the reducing agent are applied separately. First, the samples are immersed in buffered 2% OsO_4_ solution for 90 min. Subsequently, the samples are placed for 90 min in 2.5% buffered K_4_[Fe(CN)_6_] without any washing step in-between. We repeated this procedure for 4-mm brain punches: First, the samples were incubated in 2% buffered OsO_4_ for 22 hours resulting in homogeneous staining throughout the sample. Next, we placed the sample directly without any washing step in 2.5% buffered K_4_[Fe(CN)_6_] solution. Ref. (Hua, Laserstein, and Helmstaedter 2015) hypothesized that the main effect of reducing agents such as K_4_[Fe(CN)_6_] is to convert VIII-oxidized water-soluble osmium into an VI-oxidized water-soluble form which is thought to generate additional, non-polar OsO_2_ to be deposited in the membrane increasing the heavy metal content and the contrast in electron microscopy. However, we found that the heavy metal density in the K_4_[Fe(CN)_6_] immersed samples did not increase, but rather decrease from the sample surface towards the center as time progressed (Figure 2a, Supplementary Figure 5). This suggests that K_4_[Fe(CN)_6_] removes or “washes out” osmium from the sample(Litman and Barrnett 1972). This “washing effect” is stronger for longer incubation times as well as close to the sample surface and around blood vessels (anecdotal observation in electron microscopy images, data not shown). For incubation times longer than 12 hrs, K_4_[Fe(CN)_6_] readily dissolves and disintegrates the tissue (Supplementary Figure 6).

**Figure 2:**
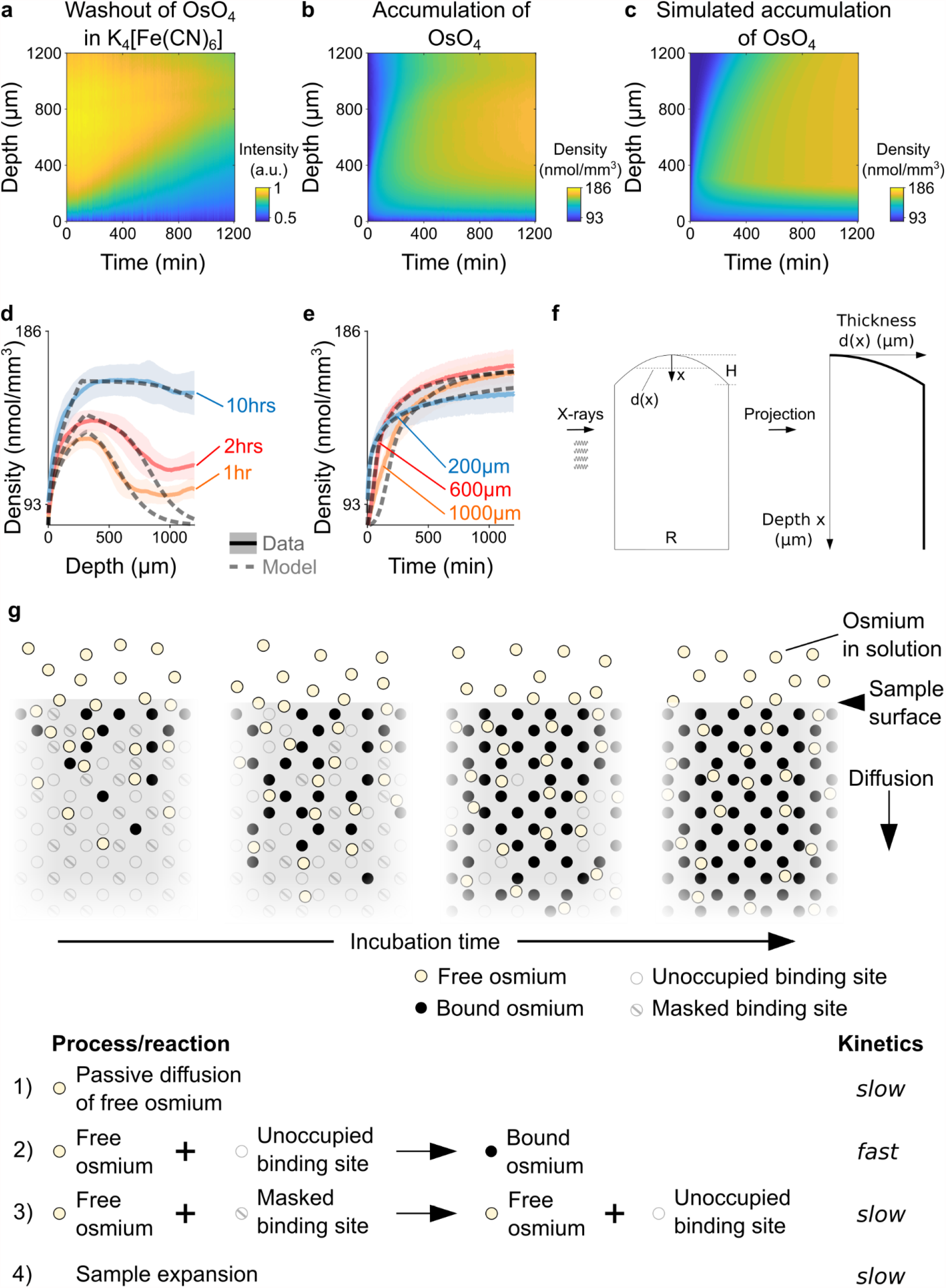
Kinetics and tissue mechanics of electron microscopy staining. **a** Spatio-temporal washout of osmium in samples incubated in 2.5% potassium ferrocyanide K_4_[Fe(CN)_6_] (n=6). The samples have been stained with 2% OsO_4_ for 22 hrs prior to placing them in K_4_[Fe(CN)_6_]. The osmium gets washed out from the sample surface towards the center as the incubation time increases. **b** Average experimentally measured spatio-temporal osmium density (nmol/mm^3^) accumulation in samples (n=13) immersed in 2% buffered OsO_4_ solution. Because of the surface curvature of the cortical samples, the tissue thickness decreases towards the surface (x=0) and results in a lower intensity in the projection images (see tissue geometry model in f). **c** Simulated spatio-temporal osmium density (nmol/mm^3^) accumulation fitted to the average experimental data in b. **d** Density profiles (nmol/mm^3^) of the spatial accumulation of osmium (n=13, mean ± s.d.) after 1 hr (orange), 2 hrs (red) and 10 hours (blue) of incubation in 2% buffered OsO_4_ overlaid with the simulation of the diffusion-reaction-advection model (dashed lines). **e** Density profiles (nmol/mm^3^) of the temporal accumulation of osmium (n=13, mean ± s.d.) at a depth of 200 μm (blue), 600 μm (red) and 1000 μm (orange) during incubation in 2% buffered OsO_4_, overlaid with the simulation of the diffusion-reaction-advection model (dashed lines). **f** The geometry of the 4-mm brain punches was modeled as a cylinder with a curved surface. The curvature of the sample surface is approximated by a circle with radius 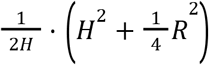, where *R* is diameter of the cylinder and *H* is the height of the curved surface. The projected tissue thickness as a function of the depth x from the sample surface is then given by 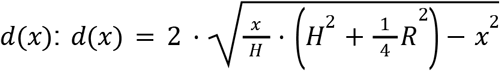, for 0≤*x*≤*H* and *d*(*x*) = *R*, for *x* > *H* **g** Diffusion-reaction-advection model for osmium staining. The model combines four coupled processes: 1) Free OsO_4_ diffuses passively into the tissue and 2) binds to the available binding sites. 3) In the presence of freely diffusing OsO_4_ previously masked binding sites are slowly turned into additional binding sites for OsO_4_. 4) The sample expands by about 5% in sample height within 20 hours of incubation (see Figure 1f).

### The kinetics of osmium tetroxide staining

The chemistry and diffusion-reaction-advection kinetics of OsO_4_ staining in aldehyde-fixed tissue are not well understood. In brain tissue, OsO_4_ is thought to mainly react with carbon-carbon double bonds, in particular of unsaturated lipids, but it has also been reported to react with some amino acids and nucleotides(Khan, Riemersma, and Booij 1961; Korn 1967; White et al. 1976; Schroeder 1980). At the beginning of the OsO_4_ incubation, the density of OsO_4_ steeply increases and peaks close to the surface of the sample. As the incubation time progresses, the OsO_4_ continues to diffuse and accumulates deeper in the tissue (Figure 2b,d,e). Due to the curved surface of the cortical tissue sample, the projected tissue thickness is lower (Figure 2f) and there are fewer binding sites for osmium, which results in a lower density of osmium closer to the sample surface(Figure 2b). At any given time, the accumulation seems to plateau from the surface towards the center of the sample, indicating that the binding of OsO_4_ saturates (Figure 2d). However, we found that the saturated density in the samples slowly increased over time from about 129 nmol/mm^3^ after 1 hour of incubation to 160 nmol/mm^3^ after 10 hours of incubation (Figure 2b,d,e), while the density of the solution remained about constant at 82 nmol/mm^3^. This suggests that additional binding sites for OsO_4_ become available slowly over time. The reactions underlying the ‘unmasking’ of binding sites are not known, but we hypothesize that it depends at least partially on the availability and the local concentration of unbound, freely diffusing osmium. Hence, at the macroscale level, the accumulation of osmium in biological tissue can be described by four coupled processes (Figure 2g): As the osmium diffuses passively into the tissue, it binds to the available binding sites in the tissue and additional binding sites are slowly being unmasked, while the tissue expands by around 5% within 20 hours of incubation (Figure 1f). The kinetics of these four processes can be described with the following coupled (non-linear) partial differential equations:

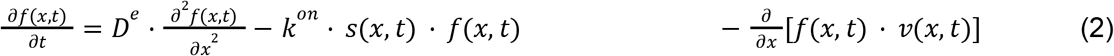

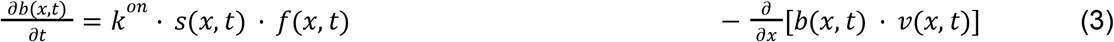

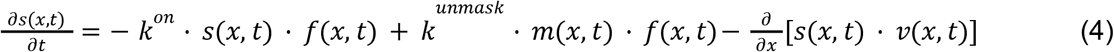

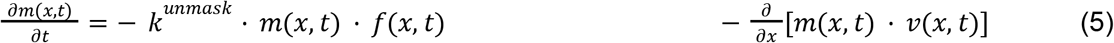

f(x,t) is the density of freely diffusing OsO_4_, b(x,t) is the density of osmium bound to the tissue, s(x,t) is the density of available binding sites and m(x,t) corresponds to the density of masked binding sites at time t and a radial depth x from the cortical surface. D^e^ is the effective diffusion coefficient for OsO_4_, k^on^ is the reaction rate constant for the binding of osmium to the tissue and k^unmask^ is the reaction rate constant at which masked binding sites get unmasked. In addition, the local change of the tissue geometry, e.g. the expansion as observed for OsO_4_, results in a velocity field v(x,t) that describes the local spatio-temporal flow of the density of each component c(x,t):

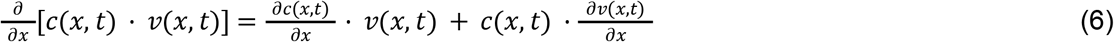

The flow consists of an advection term 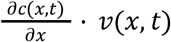, corresponding to elemental volumes moving with the local flow, and a dilution term 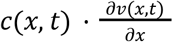 that describes the local volume change. While we assume that the tissue density is homogeneous across the cortical thickness, we consider differences in the sample thickness due to the curvature of the cortical surface. Additional information on the model assumptions can be found in the methods section.

We fitted the parameters of the system of reaction-diffusion-advection equations for a tissue sample with the projection geometry shown in Figure 2f to the average measured spatio-temporal accumulation of OsO_4_ (Figure 2b-e) as well as to the individual samples (n=13, Supplementary Figure 1). The fitted parameters of the diffusion-reaction-advection model (Supplementary Figure 7) capture the measured diffusion-reaction-advection kinetics of osmium staining well (Figure 2c, Supplementary Figure 1; residual standard error SE_res_=0.026 ± 0.005, mean ± s.d.).

## Discussion

*In situ* X-ray assisted staining is a new tool for monitoring the different steps of *en bloc* staining protocols for electron microscopy. By opening the “black box” of staining protocols, X-ray assisted staining enables, for the first time, to observe directly the diffusion of heavy metals and their microscopic effects on the tissue. While the technique will be useful to develop novel staining protocols for small to intermediate sized samples with dimensions less than 1 mm, it will particularly accelerate and facilitate the development and optimization of new staining protocols for large biological samples such as whole brains (Figure 1g).

We monitored the accumulation of osmium in different commonly used staining solutions and found significant differences in the accumulation densities. In addition, we found that the tissue expands and shrinks significantly depending on the composition of the staining solution. In particular, we found that the tissue height increased by 5% in buffered osmium solution, while the tissue height decreased by 5% in buffered reduced osmium solution. We hypothesize that this shrinkage could provide a mechanistic explanation for the long standing issue of diffusion barrier formation in large samples during reduced osmium incubation.

The method also revealed that K_4_[Fe(CN)_6_], a reducing agent that was believed to enhance the accumulation of osmium in membranes, in contrast can wash out osmium from stained tissue if applied separately. If K_4_[Fe(CN)_6_] removes osmium from the sample, why does the protocol modification of ref. (Hua, Laserstein, and Helmstaedter 2015) result in enhanced electron microscopy contrast for samples with dimensions of up to 1 mm? A potential explanation could be that K_4_[Fe(CN)_6_] clears the cytosol more efficiently than the cell membranes. As a consequence, the osmium content in the cytosol gets washed out more/faster, which might result in an overall electron microscopic contrast increase between the membranes and the cytosol. However, the chemical mechanisms by which K_4_[Fe(CN)_6_] interact with the tissue remain poorly understood and will require additional work.

X-ray assisted staining opens the door for *in silico* staining protocol optimization, by enabling to measure important heavy metal staining parameters such as the effective diffusion coefficients and reaction rate constants directly *in situ*. We derived a diffusion-reaction-advection model that describes the accumulation of osmium in homogeneous tissue by four distinct, but coupled, processes. Osmium diffuses passively into the tissue and binds there quickly to available binding sites, e.g. unsaturated lipids. Thereby the tissue expands by about 5% in height. Additional binding sites are being unmasked over the course of hours, which enables the tissue to accumulate additional osmium. While the binding of osmium appears to be nearly instantaneous, the passive diffusion and unmasking reactions are slow and appear to be rate-limiting for the staining. The mechanisms and reactions underlying the unmasking of additional binding sites for osmium are not known. Unsaturated lipids, one of the prominent binding sites for osmium, can be masked by lipid-protein complexes. In the presence of osmium, these complexes could dissociate and make the previously masked unsaturated lipid available for binding osmium(Clayton 1959; Lauder and Beynon 1993; Ashhurst 1961; Wigglesworth 1975). A recent study used time-of-flight secondary ion mass spectrometry to analyze the colocalization of different fatty acid species with osmium at the tissue surface(Belazi et al. 2009). As expected, the unsaturated fatty acids colocalize well with the tissue-bound osmium. However, they also report a remarkable complementary clustering of saturated and unsaturated fatty acids in osmium stained tissue that appears not to be present in unstained tissue. Hence, it could be that osmium can convert saturated lipids to unsaturated lipids through hydrogenation/oxidation. Alternatively, the masked binding sites could also be interpreted as a different binding partner that reacts much slower with osmium.

Biological samples such as brains are non-homogeneous media. They are composed of different tissue and cell types and therefore the diffusion of staining agents is expected to be heterogeneous. Nevertheless, the fitted model parameters reproduce the observed kinetics of osmium accumulation relatively well up to a depth of about 700 – 800 μm from the surface. Deeper in the tissue osmium that diffused from secondary sources such as ventricles or axon tracts appear to contribute to the accumulation of osmium. For example, we found that white matter and myelinated axon tracks tend to accumulate osmium faster and stronger compared to gray matter (Supplementary Figure 1-5). Such secondary sources and tissue inhomogeneities are not considered by our model. Still, the fitted model parameters appear consistent with comparable values from the literature, although we also find some discrepancies. For example, in mouse brains we expect to find about 117.975 nmol/mm^3^ double-bonds on unsaturated lipids (see methods for the derivation). Because the dominant binding reaction for osmium is assumed to be the formation of mono- and di-ester bonds with unsaturated lipids which has been described to result in the deposition of OsO_2_, the density of double bonds should roughly correspond to the density of available binding sites. But the proposed model predicts 2.999 times more binding sites, even without unmasking. One interpretation would be that unsaturated lipids are not the dominant reaction partner and that other reaction partners such particular amino acids or nucleotides account for about two third of the available binding sites. Alternatively, this could indicate that each unsaturated lipid double bond gets associated with on average 3 osmium atoms, e.g. through Os_3_O_5_^-^ and Os_3_O_5_H^-^ isotopes that have previously reported to exist in osmium-stained adipose mouse tissue(Belazi et al. 2009). Similarly, it has been reported that if the mono- and di-ester products are left standing in solution, a highly insoluble trimer osmium product is forming(Schroeder 1980).

We expect that X-ray assisted staining will be very useful in establishing a more refined picture of the heterogeneity of the heavy metal diffusion processes in different tissue types and compartments as well as in different conditions. In particular, it will be interesting to examine how changes in the extracellular space and different fixation protocols or microwave-assisted incubation affect the diffusion dynamics(Korogod, Petersen, and Knott 2015; Fulton and Briggman 2020; Jensen and Harris 1989; Login and Dvorak 1988).

Furthermore, X-ray assisted staining can be used at higher resolution and in combination with computed tomography for producing a more detailed picture of the tissue mechanics involved in different staining steps. We and several other labs have noticed that brain tissue expands significantly during incubation in “osmium amplifiers” such as the osmiophilic thiocarbohydrazide (TCH)(Seligman, Wasserkrug, and Hanker 1966). Ref. (Mikula and Denk 2015) reported that nitrogen bubbles are formed during TCH incubation that could be responsible for the formation of tissue cracks in large samples. X-ray assisted staining could be used to detect and monitor the formation of tissue cracks and to explore ways to reduce these artefacts in large samples.

Aside from accelerating the development of novel staining protocols this new method will also allow for improved closed-loop quality control of the staining process. Large samples tend to be biologically more variable and more difficult to precisely fine-tune all the parameters in the staining protocol ahead of time to account for these individual differences. Hence, X-ray assisted staining can be used to adjust the parameters in real time to compensate for these individual differences. This is particularly important for experiments in which the sample yield is low or the individual sample is precious, as for example in the case of human brains or functionally imaged brains of behaving animals that require several months of behavioral training.

## Methods

### Sample preparation

Animal use procedures were approved by the Princeton University Institutional Animal Care and Use Committee and carried out in accordance with National Institutes of Health standards. The experiments were carried out with C57BL/6J mice (Jackson Laboratories, Bar Harbor, Maine, USA) of both sexes, age ranging from 13 to 58 weeks (mean±sd: 27±9 weeks). Prior to tissue sample extraction, the mice were anesthetized by isoflurane inhalation (4%) and euthanized with an intraperitoneal injection of a ketamine (100mg/kg) and xylazine (10 mg/kg) overdose. Next, the animals were perfused transcardially with ≥ 50ml fixative solution (1.3 – 1.5% glutaraldehyde (GA) and 2.5 – 2.6% paraformaldehyde (PFA) in 0.14 – 0.15M cacodylate buffer with 2.0 – 2.1mM CaCl_2_ at pH 7.4; (GA, PFA and cacodylate buffer Electron Microscopy Sciences, Hatfield, PA; CaCl_2_, Sigma Aldrich, St. Louis, MO). After perfusion, the animal was left for 15 – 60 minutes on ice. Subsequently, the brain was removed in order to extract tissue samples at the center of the cortical surface of each hemisphere using biopsy punches of 4 mm diameter. The samples were kept in fixative solution at 4 °C for 36 hours (4-mm biopsy punches) or for 12 hours (whole mouse brain). Next, the samples were washed 6 – 7 times for 30 minutes in 0.15 M cacodylate buffer with 2 mM CaCl_2_.

Glass vials were prepared prior to the experiments with a 2-mm layer of Sylgard 184 (Electron Microscopy Sciences, Hatfield, PA) at the bottom. The whole mouse brain was glued with a small drop of Vetbond tissue adhesive (3M, Hanna Pharmaceutical Supply Co., Inc, Wilmington, DE) to the bottom of the vial. For the 4-mm biopsy punch samples, a hole with a diameter of 4 mm was cut out of the Sylgard layer. Next, the samples were placed into the sylgard hole and were glued to the bottom of the glass vial with a small drop of Vetbond tissue adhesive. Subsequently, the samples were immersed in fresh cacodylate buffer solution for the transfer to the X-ray microscope. At the microscope, presets (see X-ray microscopy section below) were loaded for image acquisition. After that the washing solution was replaced with the staining solution and the sample was placed into the X-ray microscope. The acquisition of X-ray images was typically started within 5.66 ± 2.35 min. (mean ± s.d.) after adding the staining solution. The following staining solutions have been assessed for this study (all buffered with 0.15 M cacodylate buffer and 2 mM CaCl_2_):

- Osmium tetroxide: 1%, 2% and 3% OsO_4_ (prepared from 2 mL 4% osmium tetroxide solution, Electron Microscopy Sciences, Hatfield, PA)
- Potassium ferrocyanide: 2.5% potassium ferrocyanide (prepared from crystalline 25 g potassium hexacyanoferrate(II) trihydrate powder, Sigma Aldrich, St. Louis, MO)
- Reduced osmium: 2% OsO_4_, 2.5% potassium ferrocyanide
- Reduced osmium in 11.25% formamide: 2% OsO_4_, 2.5% potassium ferrocyanide, 11.25% formamide (prepared from 25 mL formamide solution, Sigma Aldrich, St. Louis, MO)
- Reduced osmium in 5.56% formamide: 2% OsO_4_, 2.5% potassium ferrocyanide, 5.56% formamide

### X-ray microscopy

All X-ray microscopy experiments were performed on a Zeiss Xradia 520 Versa 3D X-ray microscope (Zeiss, Thornwood, NY). The samples were immersed in heavy metal solution in glass containers tightly sealed with Parafilm (Bemis Company, Inc, Neenah, WI) just before being placed in the recording chamber of the Xradia. Test images were acquired to check imaging parameters. For the biopsy punch samples, projections at a fixed view angle (rotation angle step size = 0°) were taken every 22 – 60 seconds resulting in about 1300 – 1350 images per sample over 22 hours of continuous acquisition. For the whole mouse brain, projections were taken every 22 – 60 seconds and after each projection the sample was rotated by 1°. All images were acquired using the image acquisition software Scout-and-Scan™ Control System (Zeiss, Thornwood, NY) with the following parameters:

- Objective: 0.4X
- Tube Voltage = 100 kV
- Output power = 7 W
- Source filter = air (no filter)
- Exposure time = 20 sec.
- Effective pixel size = 2 – 3 μm (biopsy punches) and 13 – 14 μm (whole mouse brain)

### Image preprocessing

First, the individual images and metadata of each projection of each sample were extracted from the proprietary *.txrm and *.xrm files as saved by the Xradia image acquisition software. In addition, a time stamp of the acquisition time point was extracted from the image metadata for each projection. A custom cross-correlation based image alignment procedure(Wanner et al. 2016) was used to correct for translational offsets between images of subsequent time points. The projection images were flat-field corrected to account for radial intensity variations due to the cone-shaped X-ray beam as follows:

1. For each sample, a median image *im* _*median*_ was calculated from the 2nd to 5th projection images right after acquisition onset.
2. The following quadratic 2d function was fitted to *log*(*im*_*median*_) using the built-in function *lsqcurvefit* in MATLAB:

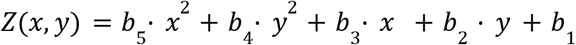
3. At each time t, the projection image was flat-field corrected using *Z*(*x, y*) with the fitted parameters:

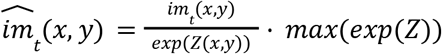

### Measuring the X-ray absorption and local heavy metal density

The pixel intensities of the detected absorbed X-rays were used as a proxy for the density of heavy metal accumulation in the tissue. In each sample, we quantified the pixel intensities in each projection taken at time t along a cross-section that pointed from the cortical surface radially towards the white matter (Figure 1b). The cross-sections were manually selected such that they started at a well detectable surface boundary and they were located at least 1 mm away from the vertical sample borders. For each sample i, the intensity *w*_*i*_ (*x, t*) of the transmitted X-rays was quantified by averaging the pixel intensity of the cross-section at a depth *x* at time *t* within a width of ±25 pixels perpendicular to the axis of the cross-section. x=0 corresponds to the cortical surface of the sample. The intensity of X-ray absorption was defined as *u*_*i*_ (*x, t*) = 10, 000 − *w* _*i*_ (*x, t*). For each time point t, the baseline intensity *b*_*i*_ (*t*) was calculated by averaging the pixels outside of the sample within -50 μm and -150 μm from the sample surface along the cross-section within a width of ±25 pixels perpendicular to the axis of the cross-section. To account for baseline fluctuation in the staining solution and the X-ray illumination the intensity of X-ray absorption was baseline-corrected by 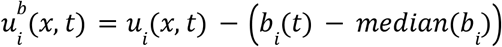.

To standardize 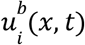, the following quantities where calculated for each sample:

1. 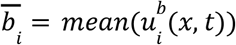 was calculated for all pixels along the cross-section outside of the sample within 0 μm and -200 μm from the sample surface.
2. 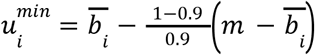
3. 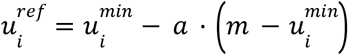
4. 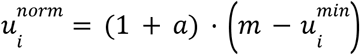 Finally, 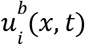 was standardized to 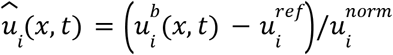

This procedure ensures that the origin is at 0 for the linear relationship between the X-ray absorption intensities in the projection images and the different OsO_4_ concentrations as shown in Figure 1c and that a pixel intensity of 0.9 roughly corresponds to the saturated staining in 2% OsO_4_. For this, the factors *a* = 0. 61749and *m* = 6420 were determined empirically.

### Measuring the linear relationship between X-ray absorption and OsO_4_ concentration

The X-ray absorption/pixel intensity of cacodylate-buffered solutions with 1%, 2% and 3% OsO_4_ was measured the same way as for the X-ray absorption in stained tissue.

### Monitoring of tissue expansion and shrinkage

Intensity based heuristics were used in a semi-automated procedure to detect the cortical surface in each image of each sample. The glass vial bottom was used as a reference point to measure the distance to the cortical surface *H*(*t*) at time *t*. Right after placing the samples in the X-ray the height of the samples was *H*_0_ = *H*(*t* = 0) and the change in sample height (expansion or shrinkage) was quantified as 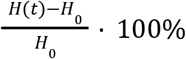.

### Tissue expansion modeling

The expansion of the tissue in buffered OsO_4_ solution was modeled by the monomolecular, saturating growth function 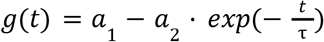. The built-in function *lsqcurvefit* of MATLAB was used for parameter fitting. The fitted parameters for the average expansion in Figure 1e were *a*_1_ = 5. 8697%,*a*_2_ = 5. 6924%,τ = 588. 23 *min*. Under the assumption that the expansion of the tissue domain in buffered osmium solution is isotropic, the corresponding velocity field v(x,t) of the flow in equation (6) can be characterized by:

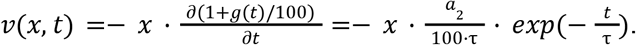

### Tissue projection geometry model

While we assumed that the tissue density is homogeneous across the brain punch, we took into account the variation in the tissue thickness of the 4-mm brain punch. Figure 2f shows a sketch of the modeled tissue geometry. The geometry of the 4-mm brain punches is modeled as a cylinder with a curved surface. The diameter of the cylinder is *R* and the curvature of the sample surface is approximated by a circle with radius 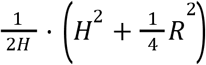. The thickness of the sample as a function of the depth x from the sample surface is then given by *d*(*x*):

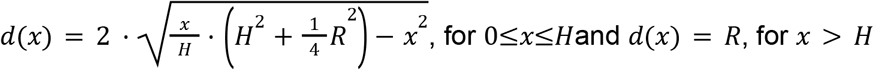

### Diffusion-reaction-advection modeling

The diffusion-reaction-advection model described by equations (2) – (5) makes the following assumptions:

– the cortical surface is in contact with an infinite source of staining agents at constant concentration,
– the composition of the 4-mm brain punches can be approximated by a semi-infinite, homogeneous medium,
– the tissue thickness is given by 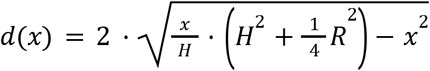, for 0≤*x*≤*H*and *d*(*x*) = *R*, for *x* > *H*
– the kinetics of OsO_4_ diffusion can be described by Fick’s second law,
– the binding of OsO_4_ to the tissue is irreversible and can be described by first-order reaction kinetics
– the density of available OsO_4_ binding sites is limited,
– the unmasking of additional binding sites is irreversible, follows first-order reaction kinetics and depends on the local concentration of free OsO_4_
– the tissue expands isotropically and can be modeled by a monomolecular, saturating growth function, which induces a local velocity field 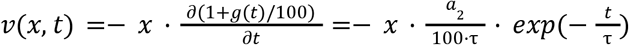.

Because the projected intensity also includes any X-ray absorbing components surrounding the sample, we have to consider the change in free osmium density outside of the sample *o*(*x, t*) for fitting the model to the experimental data:

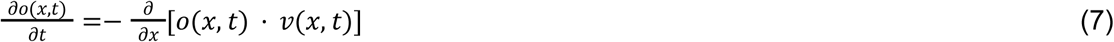

The initial conditions (t=0) for the system of coupled partial differential equations (2) – (5) and (7) were chosen as follows:

– concentration of free OsO_4_ in the sample: *f*(*x* ≤ 0, *t* = 0) = *C*^0^ · *R* and *f*(*x* > 0, *t* = 0) = 0
– density of bound OsO_4_: *b*(*x, t* = 0) = 0
– density of available binding sites: *s*(*x* ≤ 0, *t* = 0) = 0 and *s*(*x* > 0, *t* = 0) = *S*^0^ · *d*(*x*)
– density of masked binding sites: *m*(*x* ≤ 0, *t* = 0) = 0 and *m*(*x* > 0, *t* = 0) = *M* ^0^· *d*(*x*)
– concentration of free OsO_4_ surrounding the sample: *o*(*x* ≤ 0, *t* = 0) = *C*^0^ · (*D* − *R*) and *o*(*x* > 0, *t* = 0) = *C*^0^ · (*D* − *d*(*x*)), where *D* = 25*mm* is the vial diameter.

The boundary conditions for the system of coupled partial differential equations (2) – (5) and (7) were chosen within for *x* ∈ [0, *L*]with *L* = 3*mm* as follows:

– concentration of free OsO_4_ : *f*(*x* = 0, *t*) = *C*^0^ · *R* and *f*(*x* = *L, t*) = 0
– density of bound OsO_4_ : *b*(*x* = 0, *t*) = 0 and *b*(*x* = *L, t*) = *B*^0^ · *d*(*L*)
– density of available binding sites: *s*(*x* = 0, *t*) = 0 and *s*(*x* = *L, t*) = *S*^0^ · *d*(*L*)
– density of masked binding sites: *m*(*x* = 0, *t*) = 0 and *m*(*x* = *L, t*) = *M*^0^ · *d*(*L*)
– concentration of free OsO_4_ surrounding the sample: *o*(*x* = 0, *t*) = *C* ^0^· (*D* − *R*) and *o*(*x* = *L, t*) = *C*^0^ · (*D* − *d*(*L*))
– No flux beyond 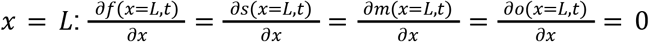

The MATLAB function *pdepe* was used to solve/simulate the initial-boundary value problem described above.

### Diffusion-reaction-advection model fitting

*C*^0^ was calculated by averaging 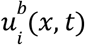 over *x* ∈ [− 150, − 50]μm from the sample surface (i.e. outside of the sample) and *t* ∈ [0, 1200]min and dividing it by the vial diameter *D* = 25*mm* and by the intensity-to-concentration scaling factor 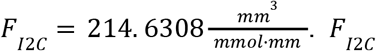 was calculated from the linear fit in figure 1c. In each fitting iteration, simulated data was produced according to the diffusion-reaction-advection model above. Within the fitting procedure, *F* _*I*2*C*_ · (*o*(*x, t*) + *f*(*x, t*) + *b*(*x, t*)) was fitted to 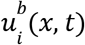 in a least square displacement sense within the domain *x* ∈ [100, 700]μm from the sample surface and *t* ∈ [0, 1200]min by using the *trust-region-reflective* algorithm of the nonlinear least-squares solver *lsqnonlin* in MATLAB to optimize the N=6 model parameters *D*^*e*^, *k*^*on*^, *k*^*unmask*^, *S*^0^, *M*^0^ and *H*. These parameters were fitted to the average experimentally measured osmium accumulation from n=13 samples within the domains *x* ∈ [100, 700]μm and *t* ∈ [0, 1200] min. To evaluate the quality of the model fits, the residual standard error was calculated as 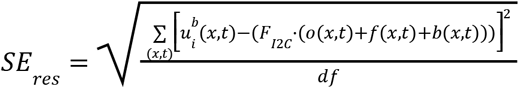, were *df* corresponds to the degree of freedom.

### Estimation of the average double-bond/ osmium binding site density in the mouse brain

The average phospholipid density in the mouse brain is 56,241±4,000 nmol/g (Barceló-Coblijn et al. 2006) and for an average mouse brain volume of 508.91±23.42 mm^3^ (Badea, Ali-Sharief, and Johnson 2007) with a weight of 0.427±0.024 g (“The Jackson Laboratory” 2021), a cubic millimeter brain tissue contains about 47.19 nmol phospholipds. Lipids in the brain carry on average 2.5 double bonds(Fitzner et al. 2020) which results in 117.975 nmol/mm^3^ double bonds to which osmium tetroxide could bind through osmium mono- or di-ester formation. Our proposed model predicts 2.999 times more binding sites (353.774±27.460 nmol/mm^3^), even without unmasking (Supplementary Figure 7).

## Acknowledgements

We thank Michael Bozlar for advice on potassium ferrocyanide and osmium chemistry, Stefan N. Oline for assistance with animal perfusions and Alyssa M. Wilson and Zhihao Zheng for comments on the manuscript. The authors acknowledge the use of Princeton’s Imaging and Analysis Center, which is partially supported through the Princeton Center for Complex Materials (PCCM), a National Science Foundation (NSF)-MRSEC program (DMR-2011750). A.A.W. acknowledges support by the CV Starr Fellowship in Neuroscience by the Princeton University. This work was supported by the National Institutes of Health (NIH) grants U19 NS104648, NIH/NEI (1R01EY027036 to HSS), NIH/NINDS (1R01NS104926 to HSS), NINDS/NIH (U01NS090562 to HSS).

## Author contributions

S.S., E.W.H. and A.A.W. designed the study and performed the experiments. A.A.W. developed the method, analyzed the data and developed the diffusion-reaction-advection models with contributions from D.W.T and H.S.S. A.A.W., S.S. and E.W.H. wrote the manuscript with contributions from all authors.

## Supplementary Information

**Supplementary Figure 1:**
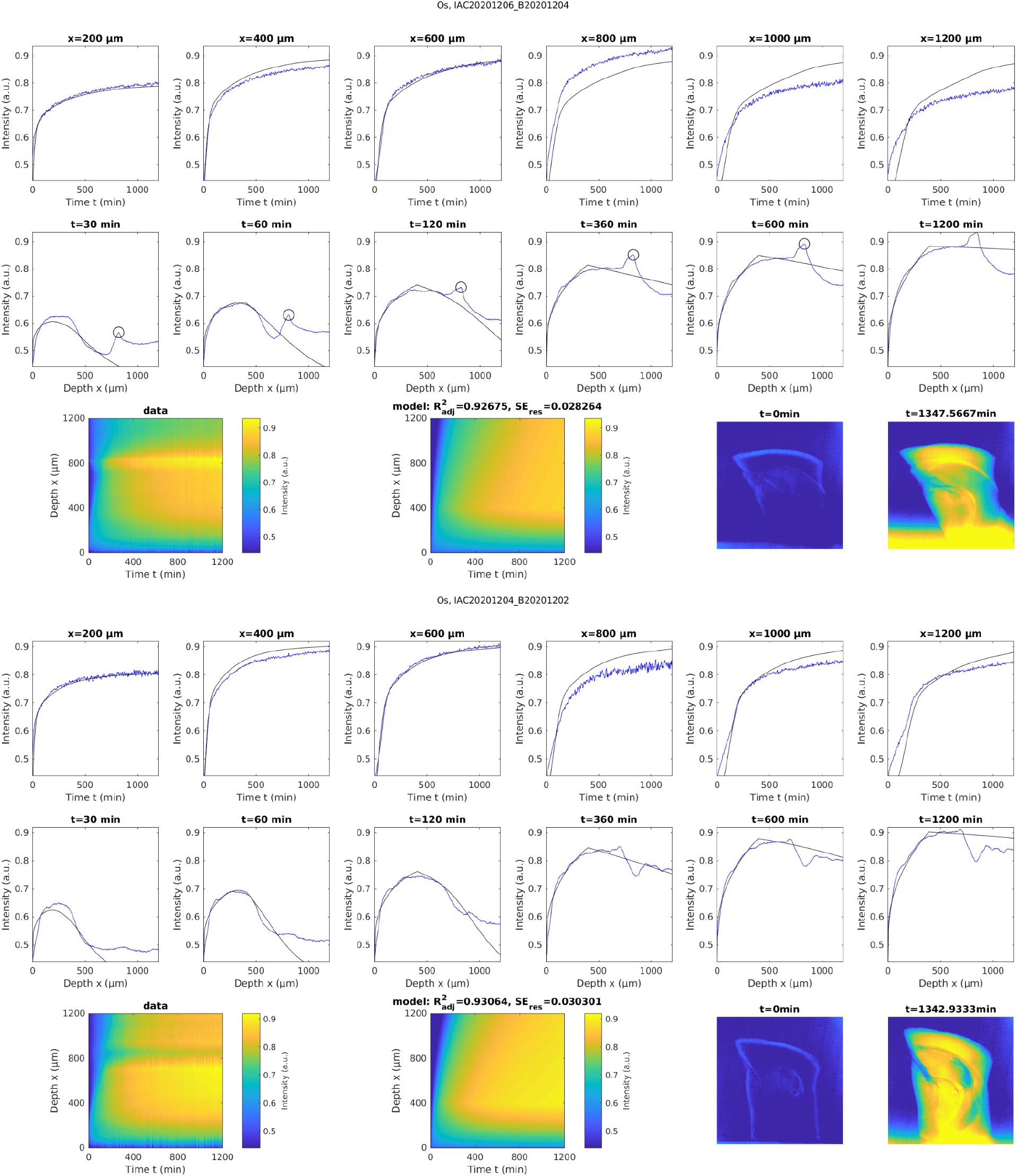
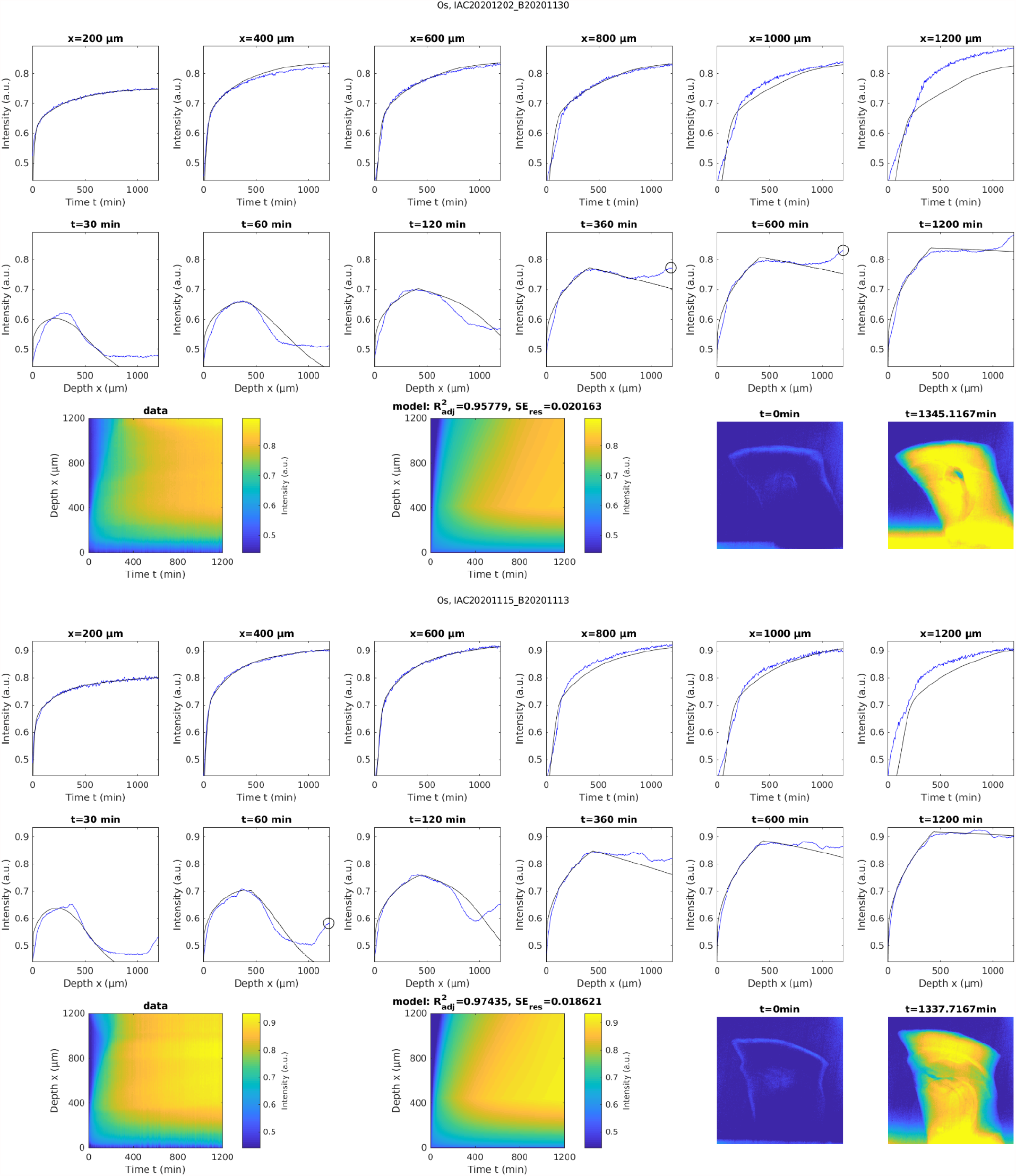
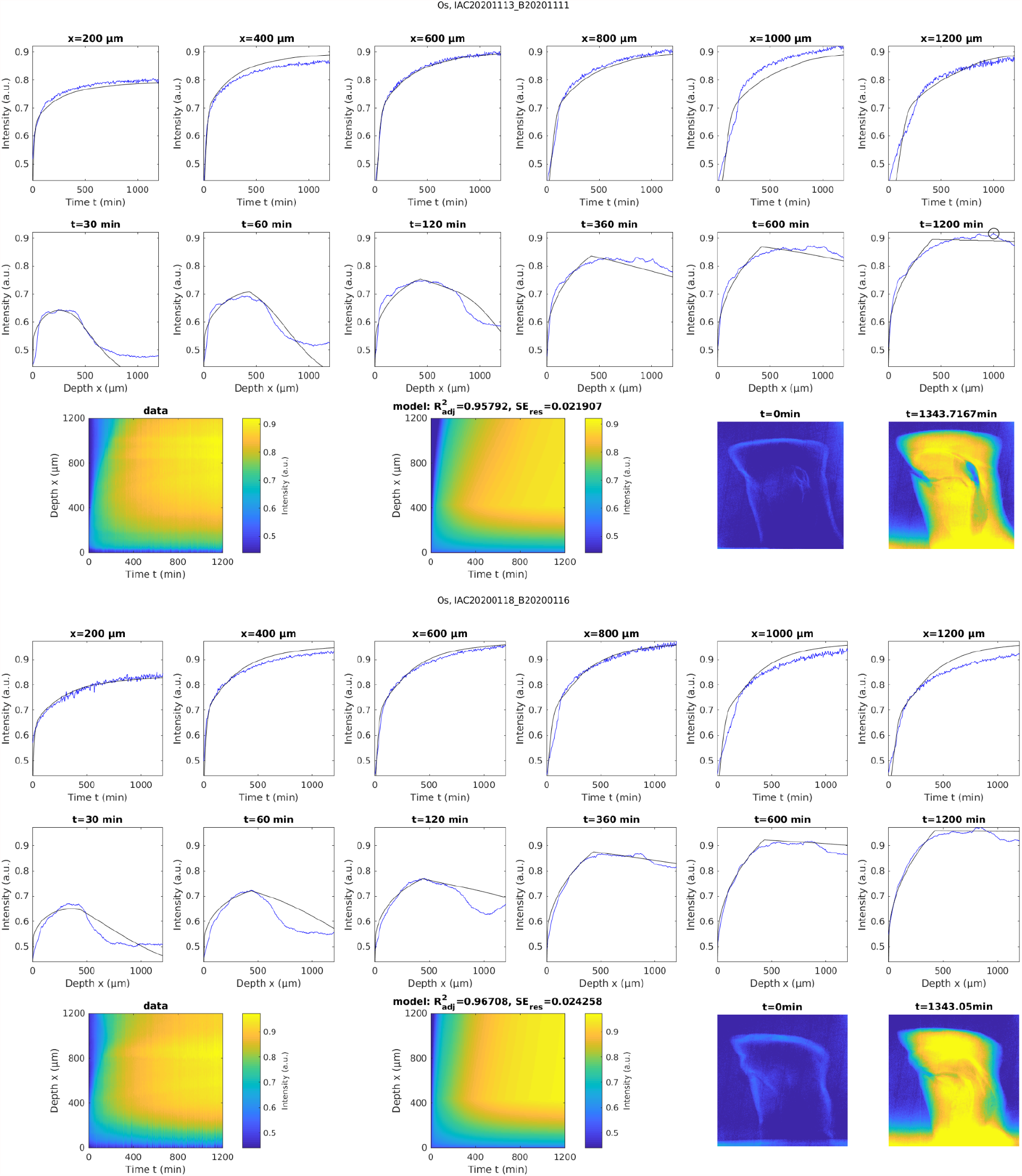
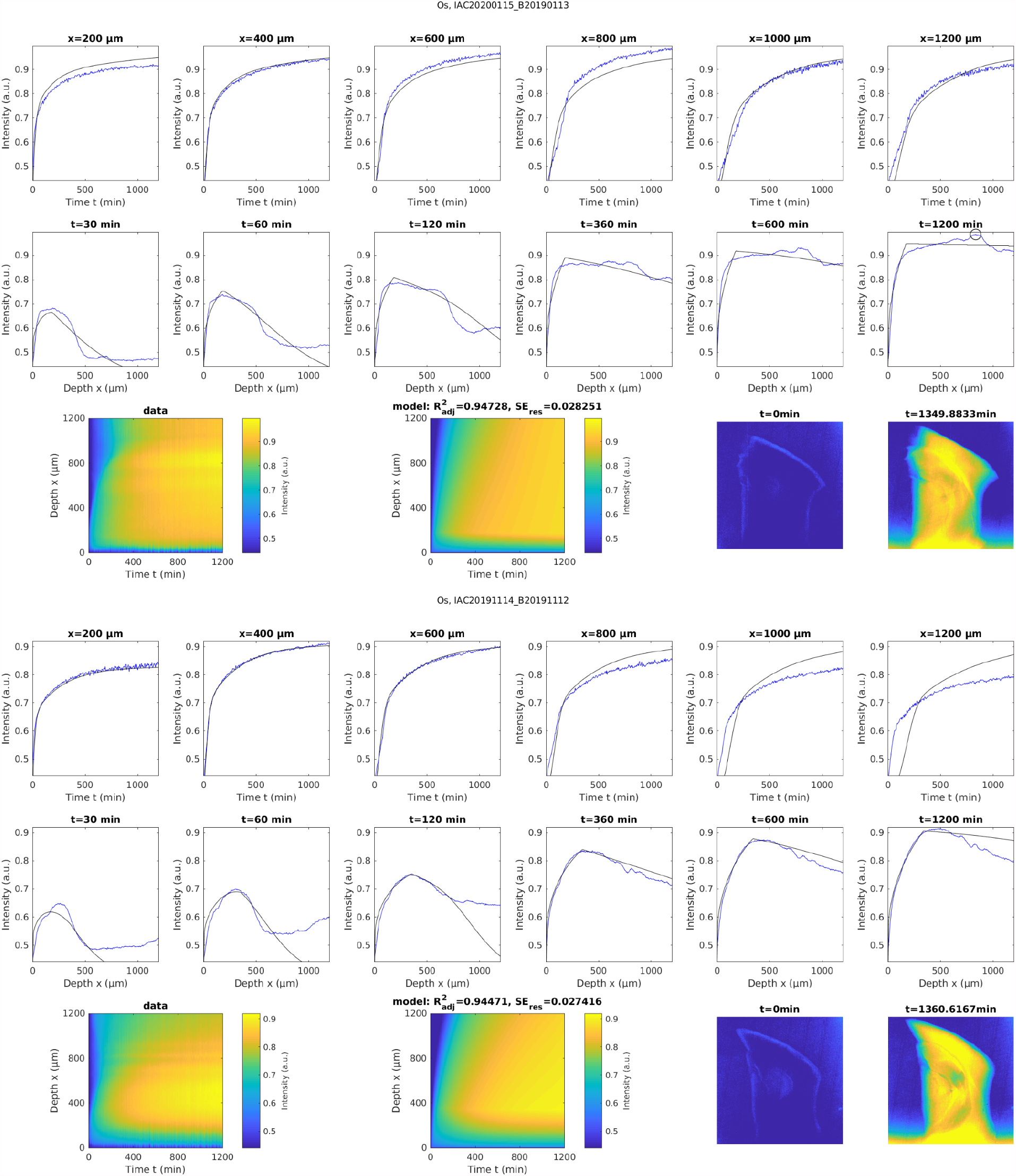
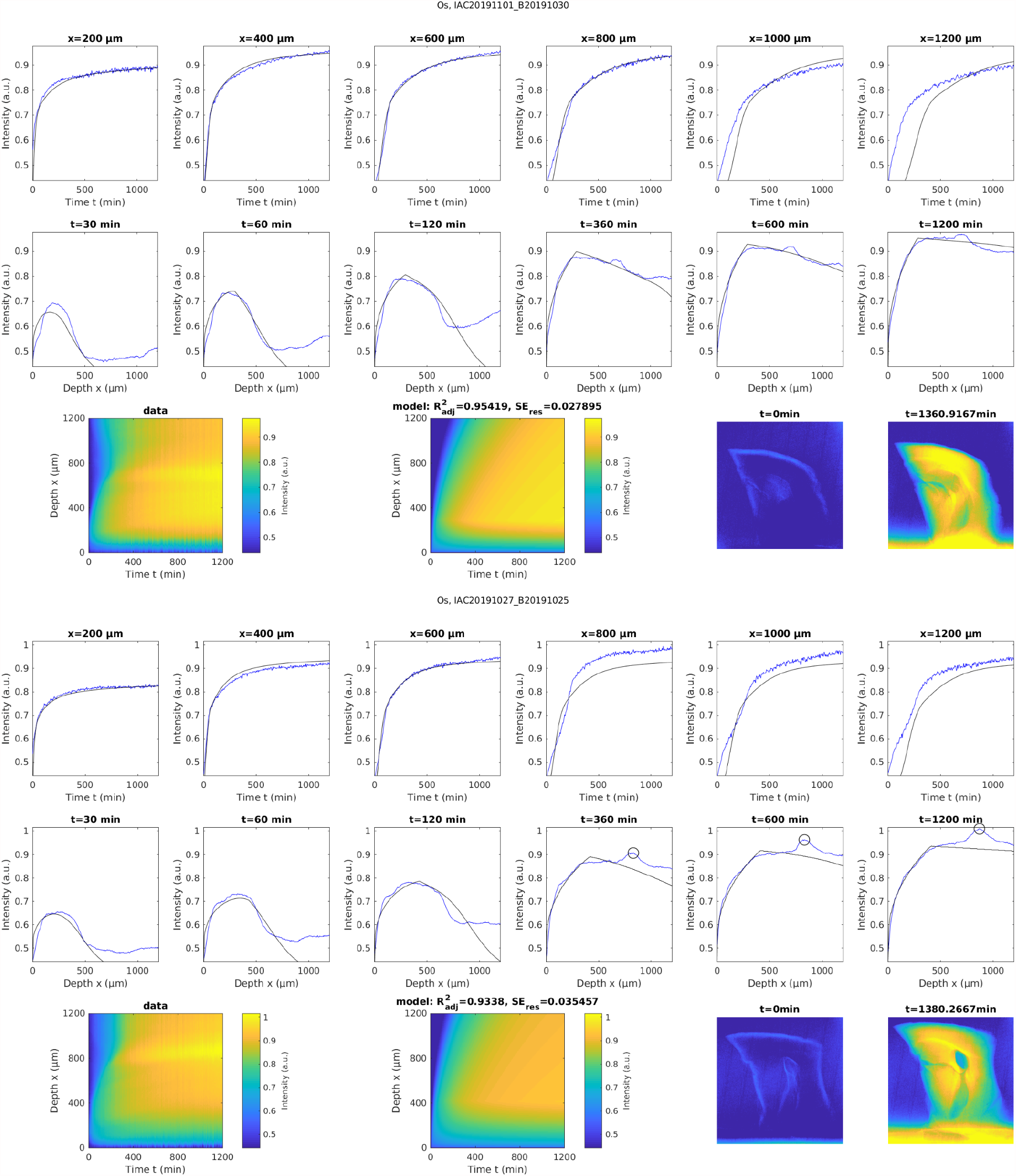
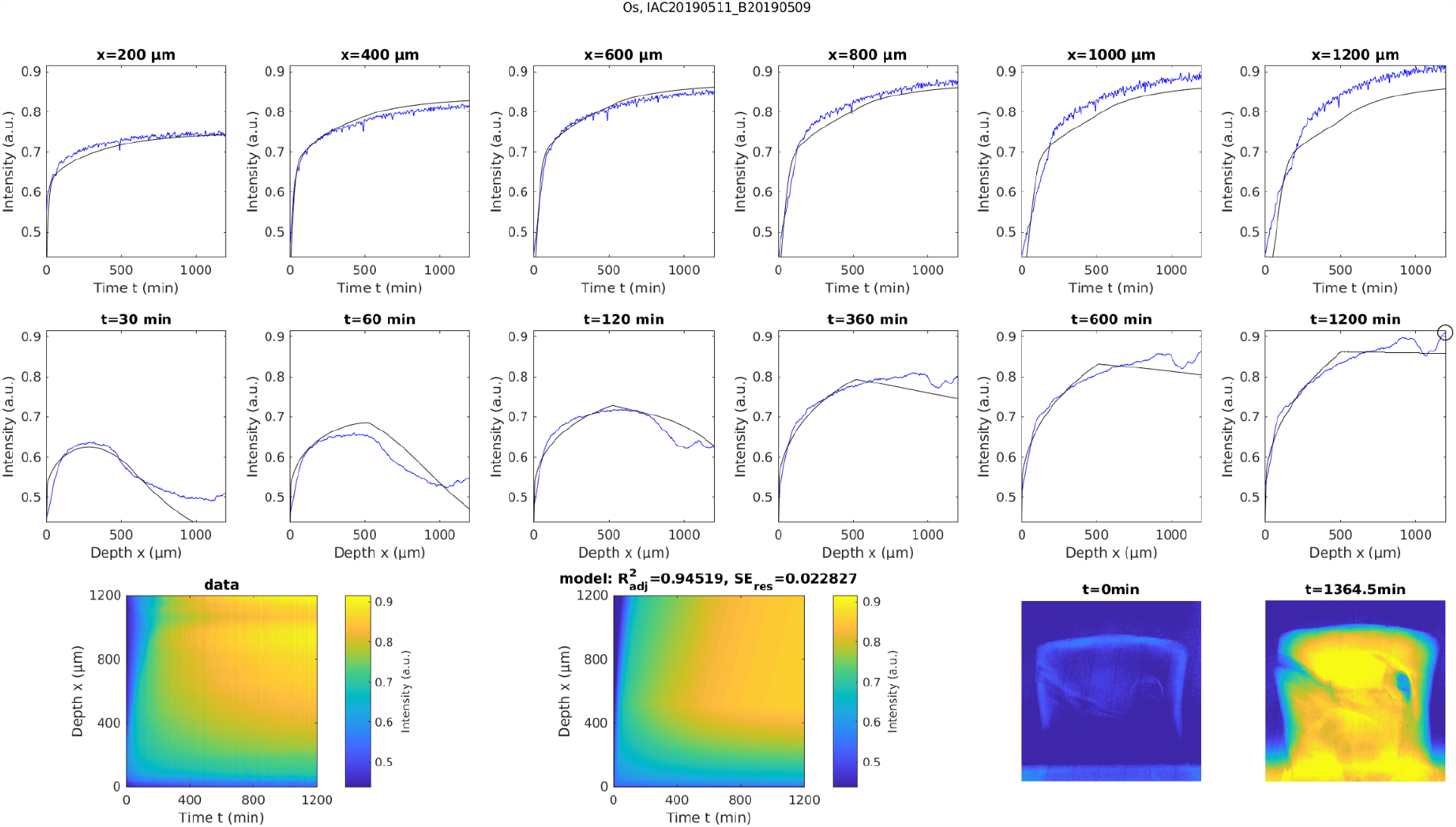
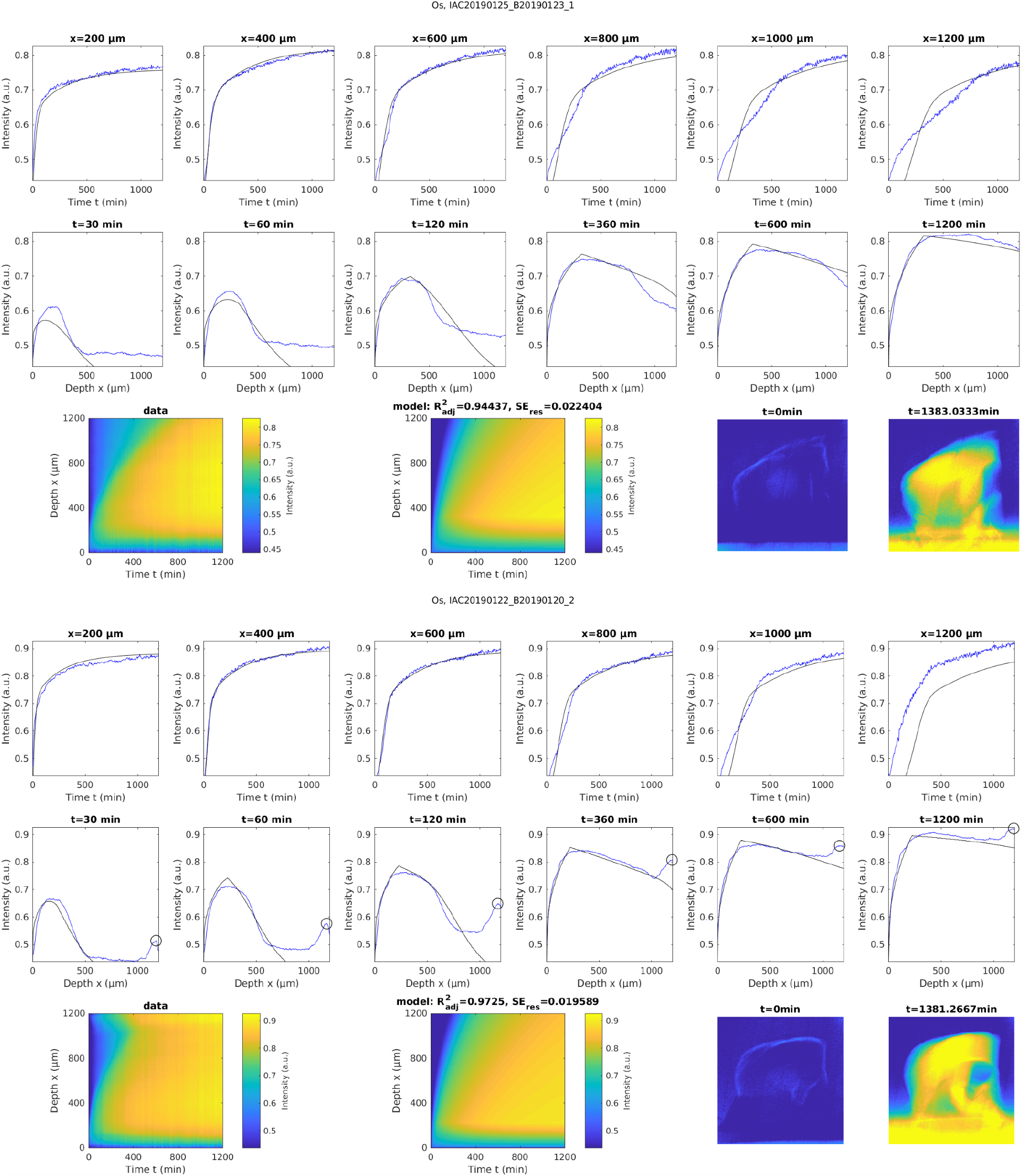
Experimentally measured accumulation of heavy metals (blue traces) in buffered 2% OsO_4_ solution in n=13 different samples. The corresponding simulated model was fitted individually to the experimental data of each sample (black traces). The first row of every sample shows the temporal profile of osmium accumulation at different cortical depths, while the second row shows the spatial profile of osmium accumulation at different times after staining onset. The black circles indicate the location of axon tracts that tended to accumulate osmium faster and stronger. The black circles indicate the location of axon tracts that tended to accumulate osmium faster and stronger. The third row shows the measured spatial-temporal accumulation of osmium (left), the corresponding fitted model simulations (middle) and the distribution of osmium in projection images of the sample at the beginning of the experiment and after about 22 hours of incubation.

**Supplementary Figure 2:**
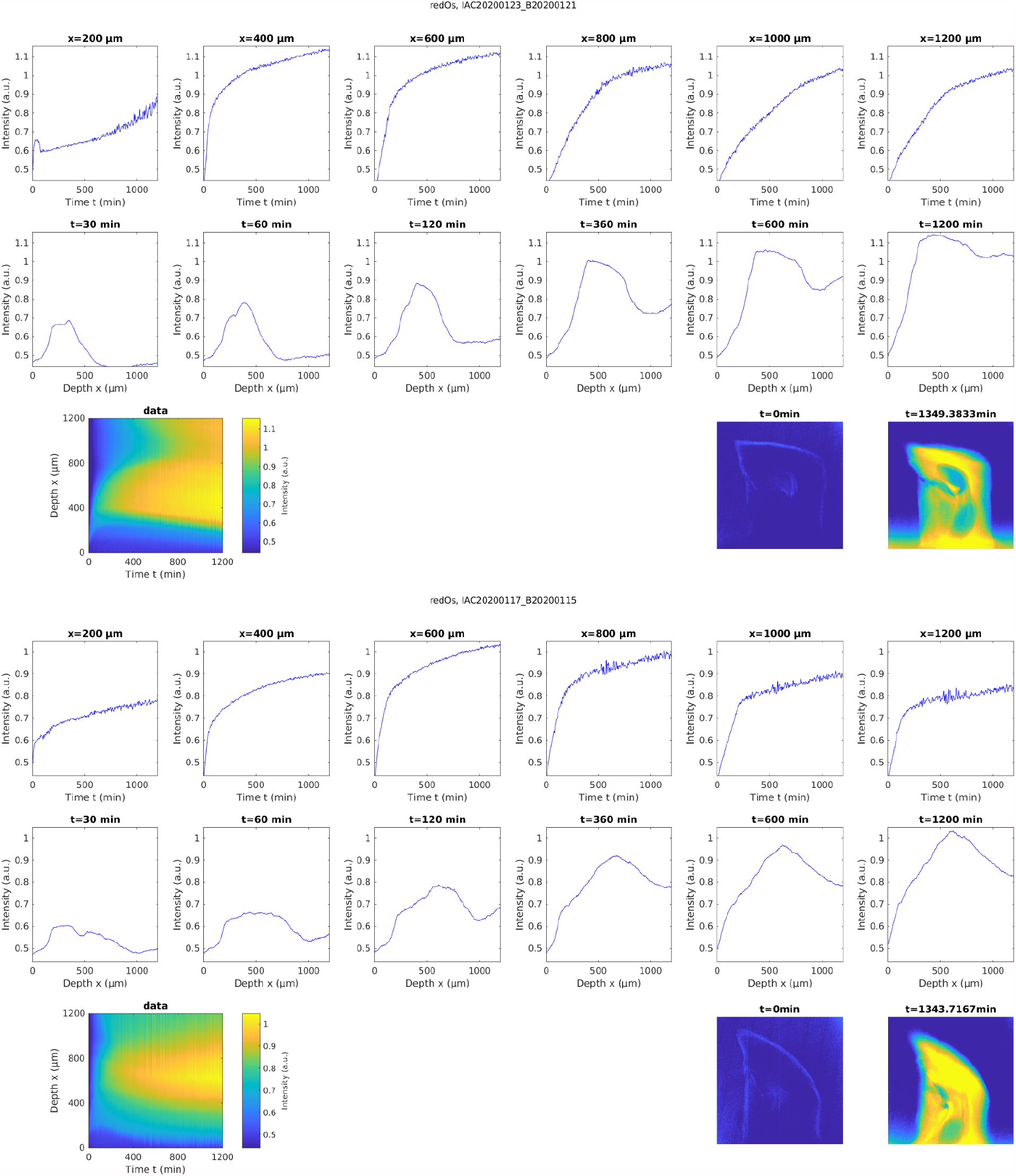
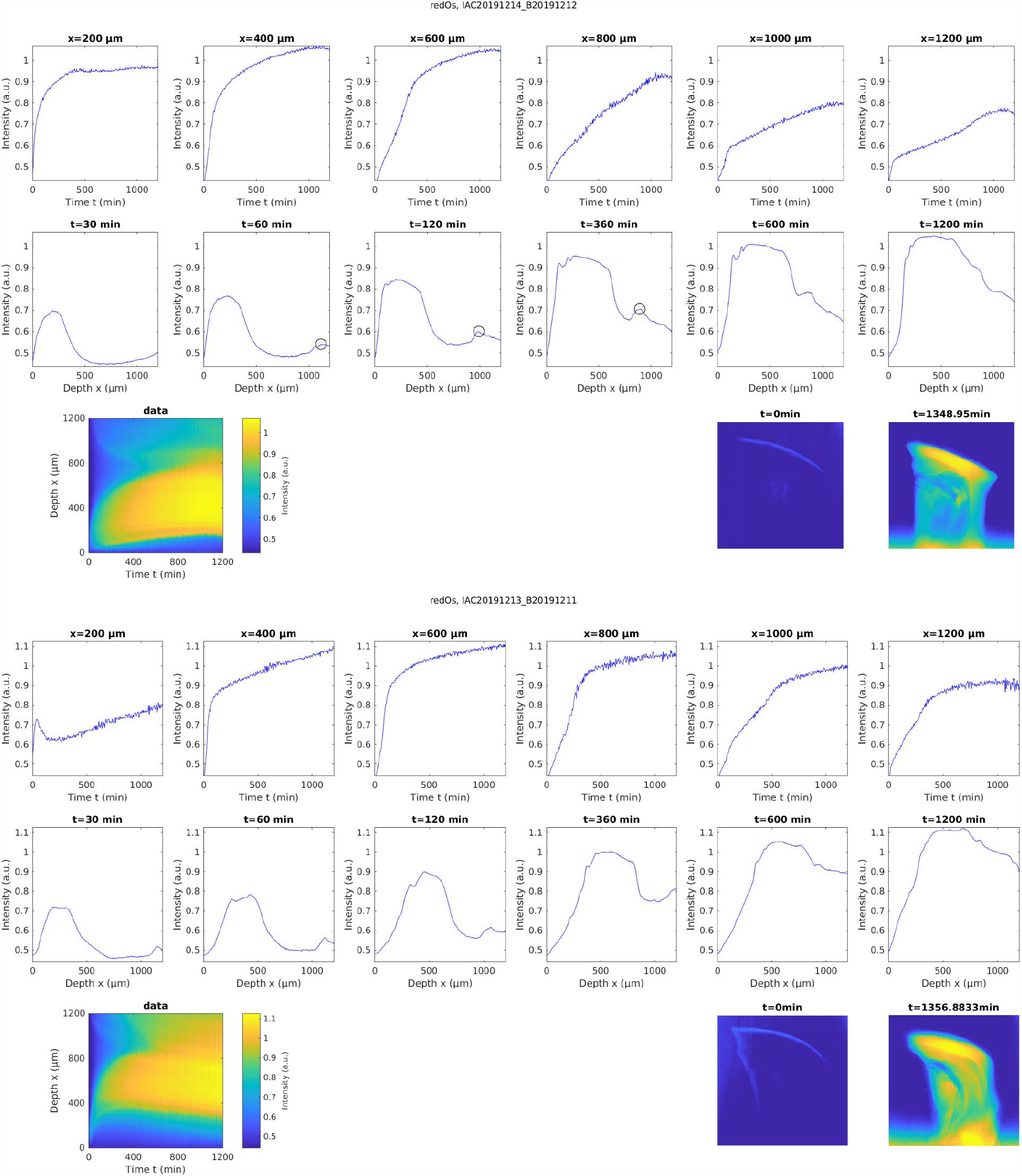
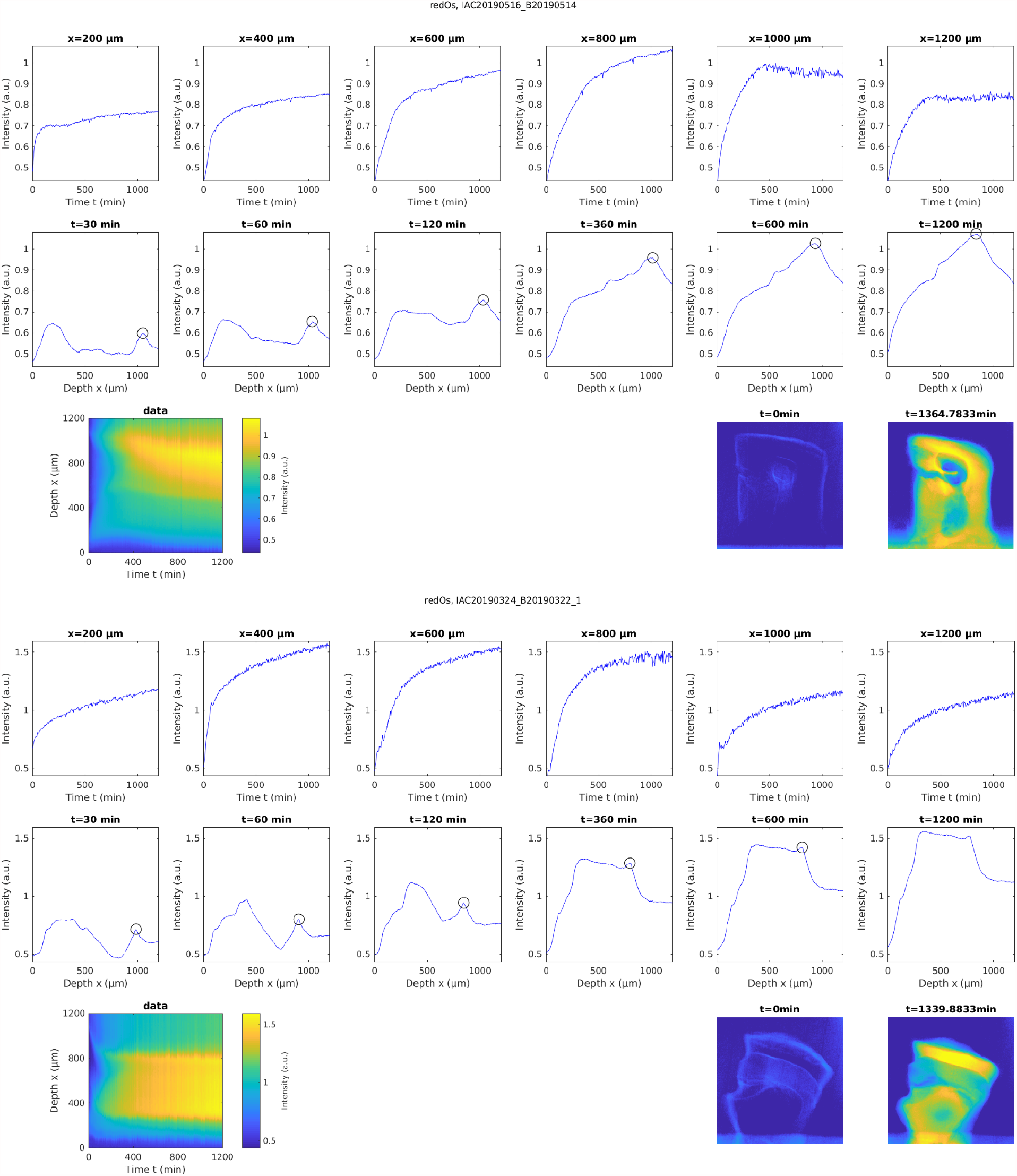
Experimentally measured accumulation of heavy metals in buffered 2% reduced OsO_4_ solution in n=6 different samples. The first row shows the temporal profile of osmium accumulation at different cortical depths, while the second row shows the spatial profile of osmium accumulation at different times after staining onset. The black circles indicate the location of axon tracts that tended to accumulate osmium faster and stronger. The third row shows the measured spatial-temporal accumulation of osmium (left) and the corresponding distribution of osmium in projection images of the sample at the beginning of the experiment and after about 22 hours of incubation.

**Supplementary Figure 3:**
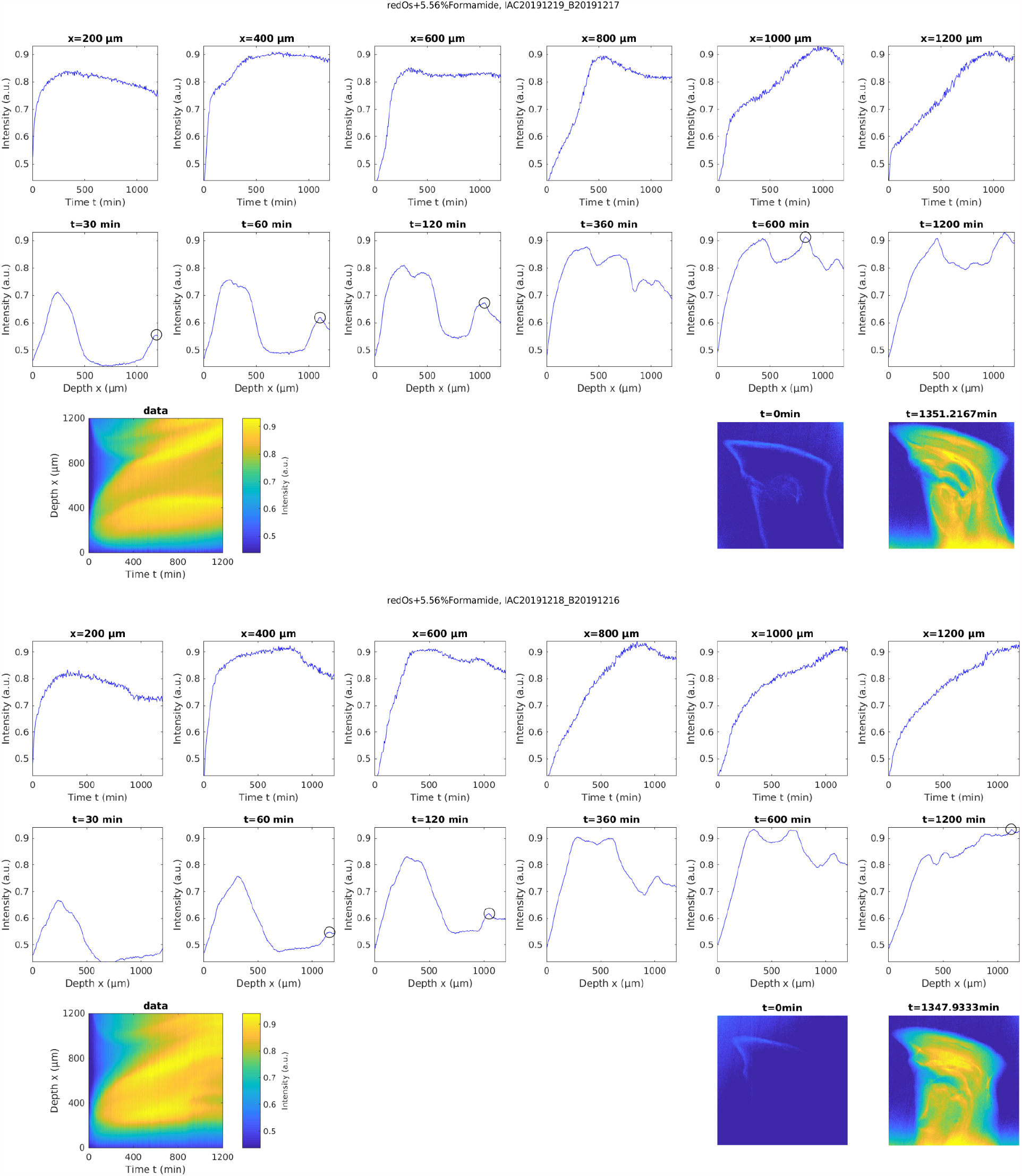
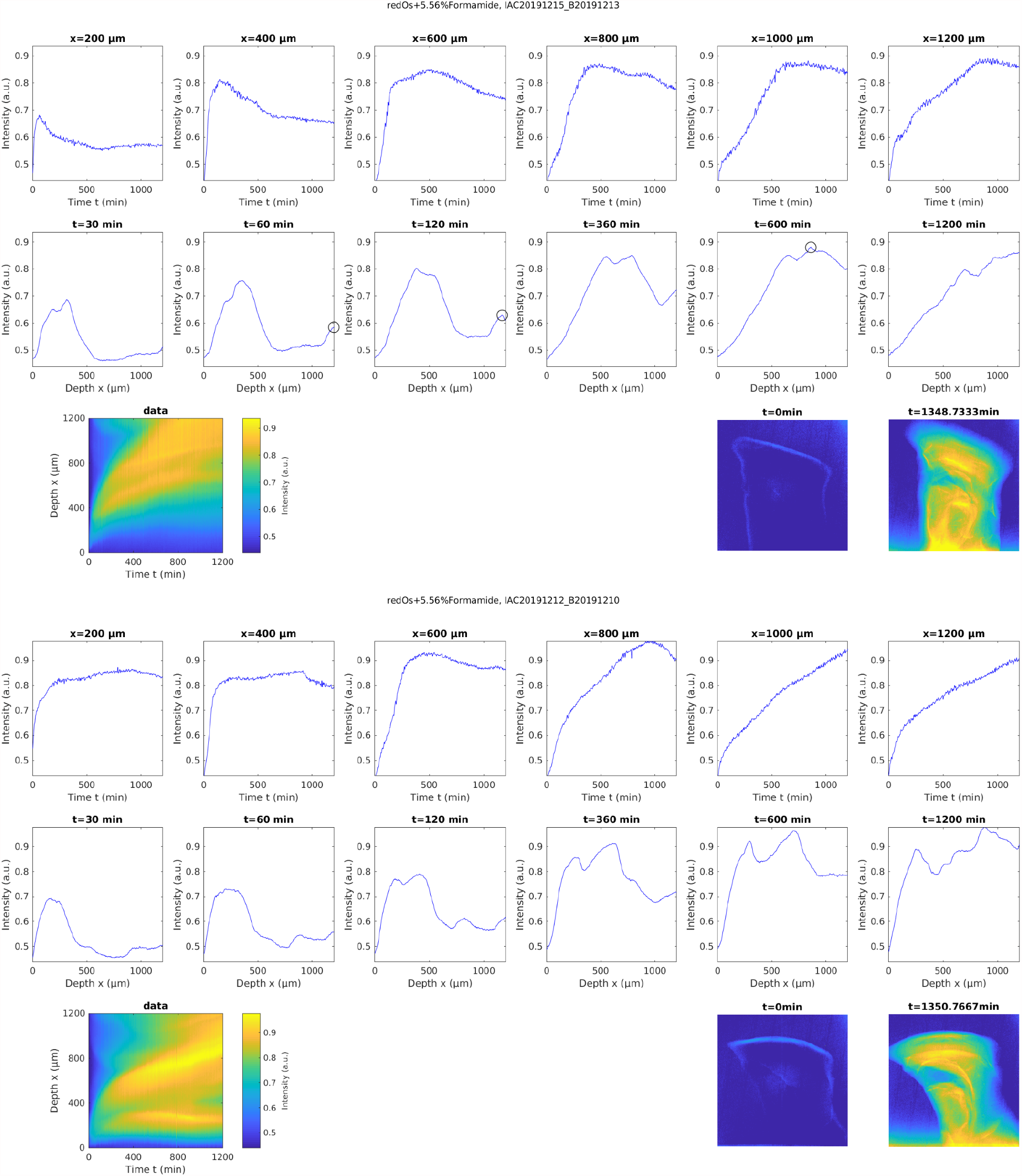
Experimentally measured accumulation of heavy metals in buffered 2% reduced osmium solution with 5.56% formamide in n=4 different samples. The first row of every sample shows the temporal profile of osmium accumulation at different cortical depths, while the second row shows the spatial profile of osmium accumulation at different times after staining onset. The black circles indicate the location of axon tracts that tended to accumulate osmium faster and stronger. The third row shows the measured spatial-temporal accumulation of osmium (left) and the corresponding distribution of osmium in projection images of the sample at the beginning of the experiment and after about 22 hours of incubation.

**Supplementary Figure 4:**
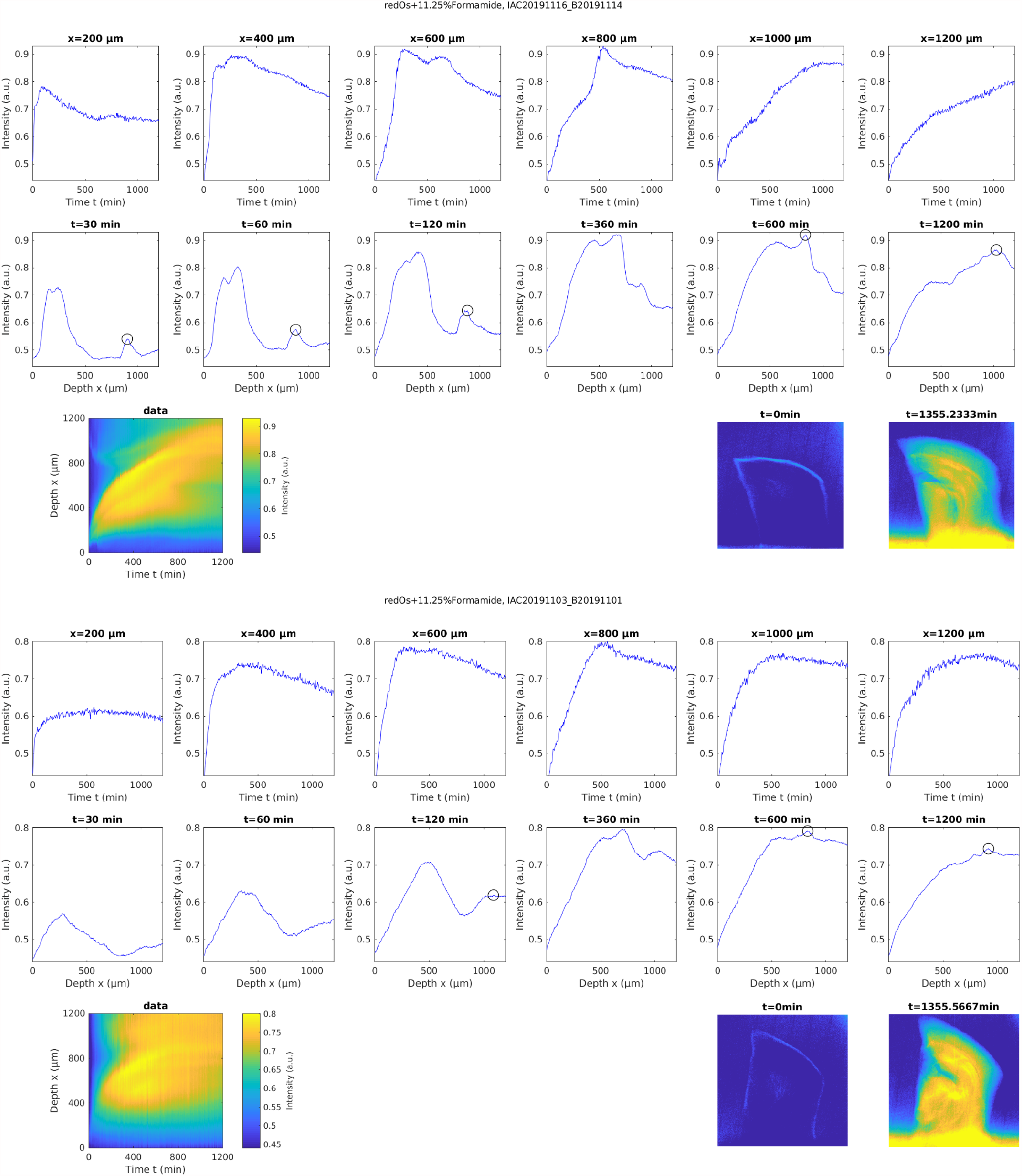
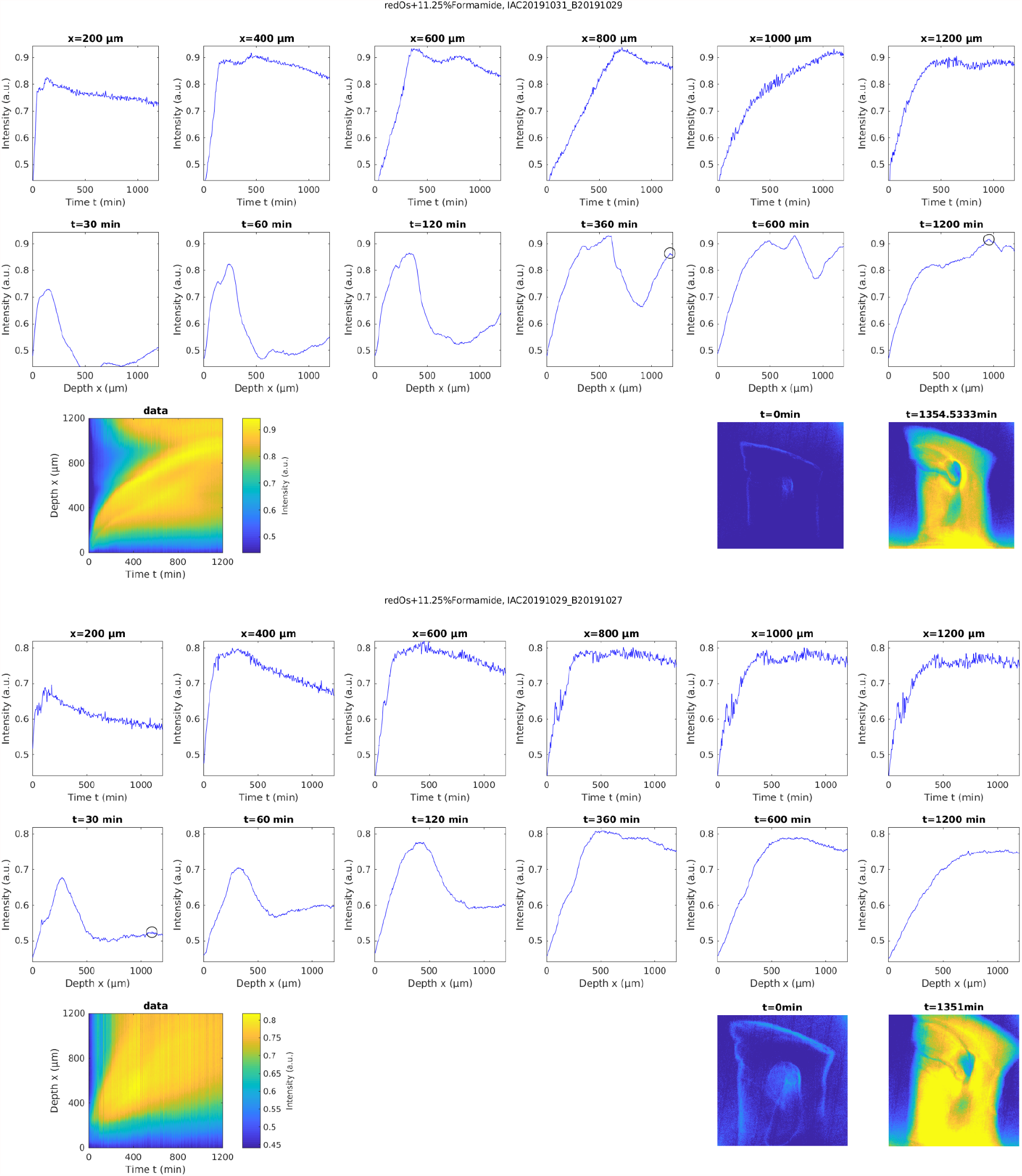
Experimentally measured accumulation of heavy metals in buffered 2% reduced osmium solution with 11.25% formamide in n=4 different samples. The first row of every sample shows the temporal profile of osmium accumulation at different cortical depths, while the second row shows the spatial profile of osmium accumulation at different times after staining onset. The black circles indicate the location of axon tracts that tended to accumulate osmium faster and stronger. The third row shows the measured spatial-temporal accumulation of osmium (left) and the corresponding distribution of osmium in projection images of the sample at the beginning of the experiment and after about 22 hours of incubation.

**Supplementary Figure 5:**
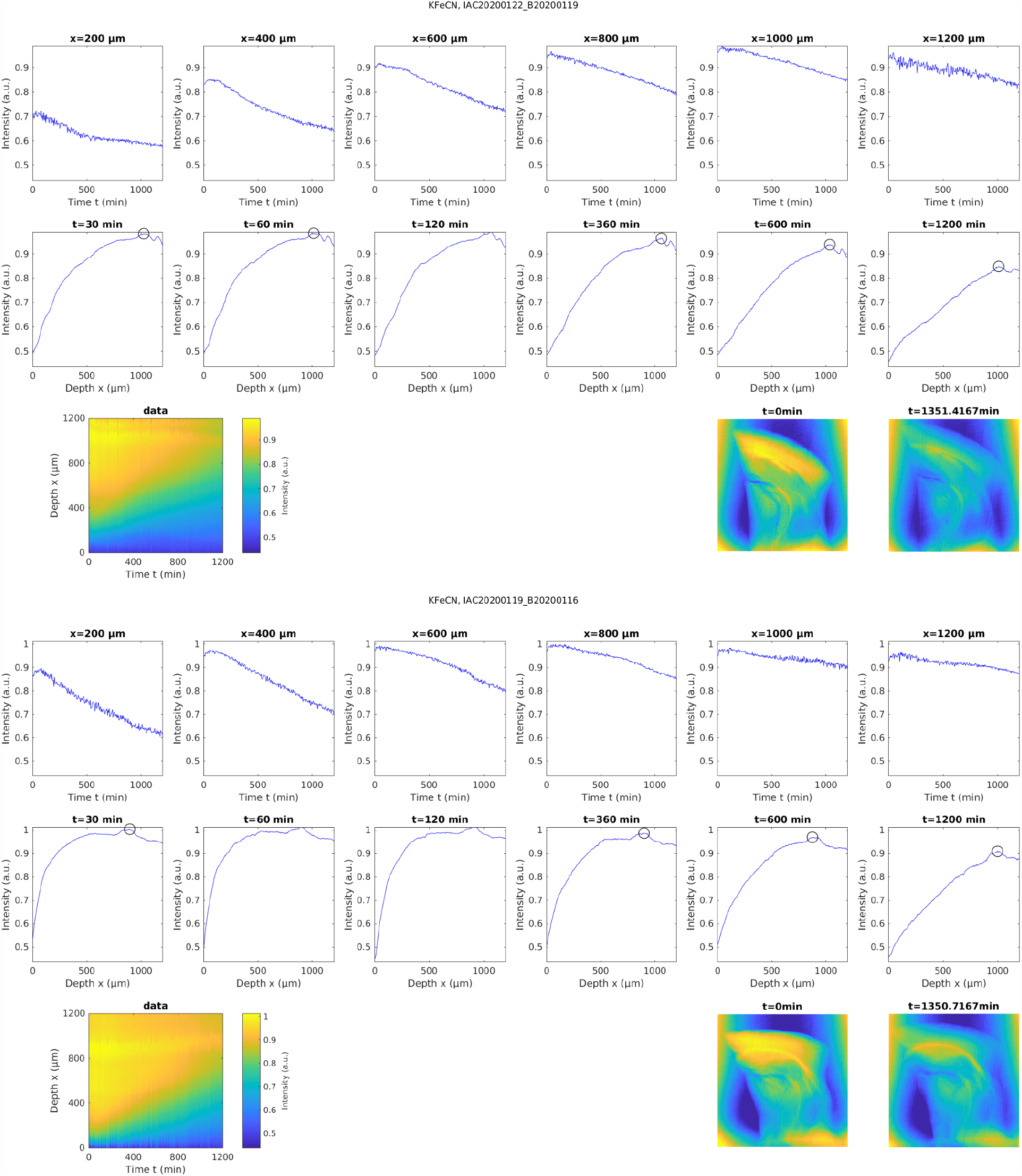
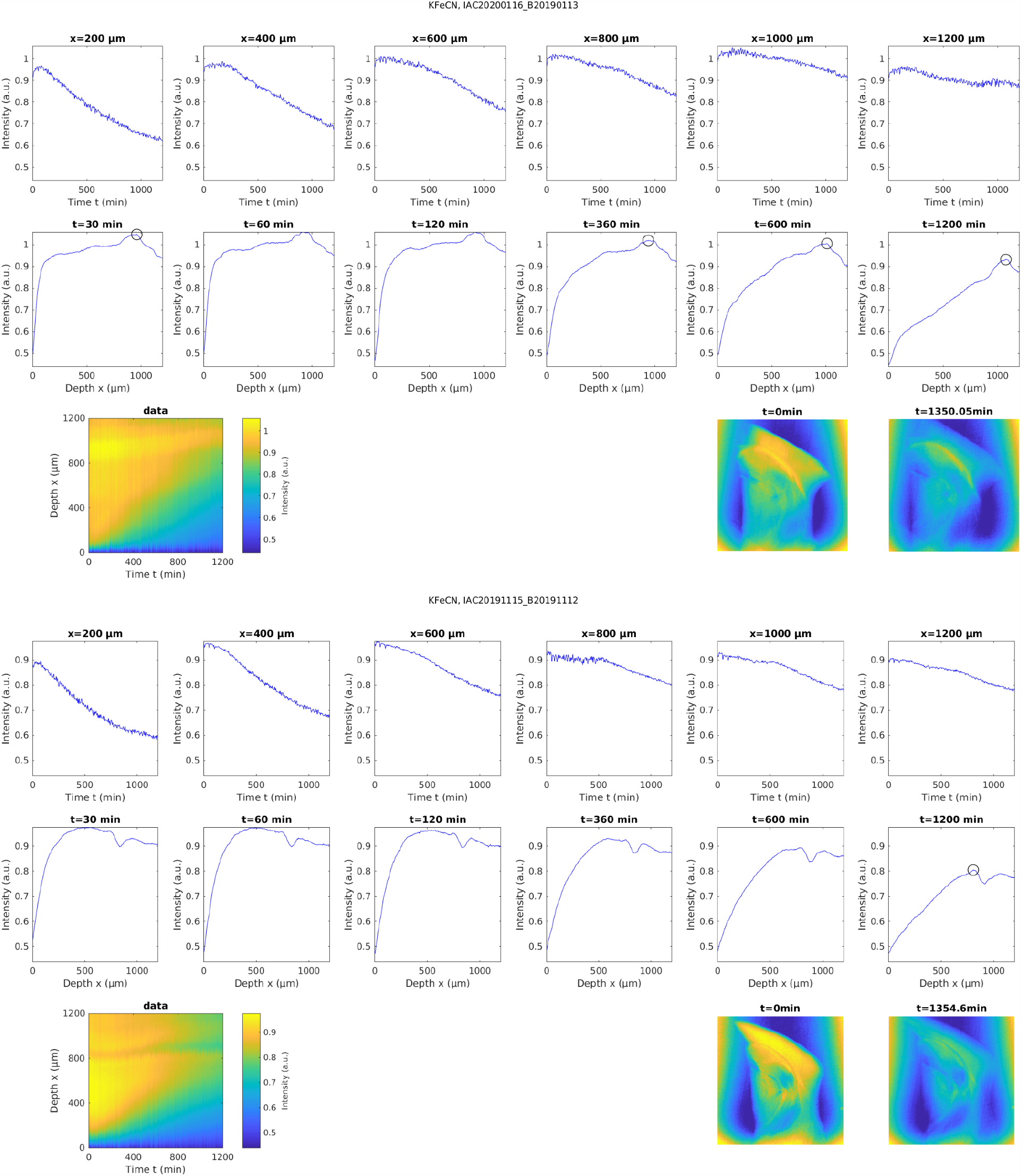
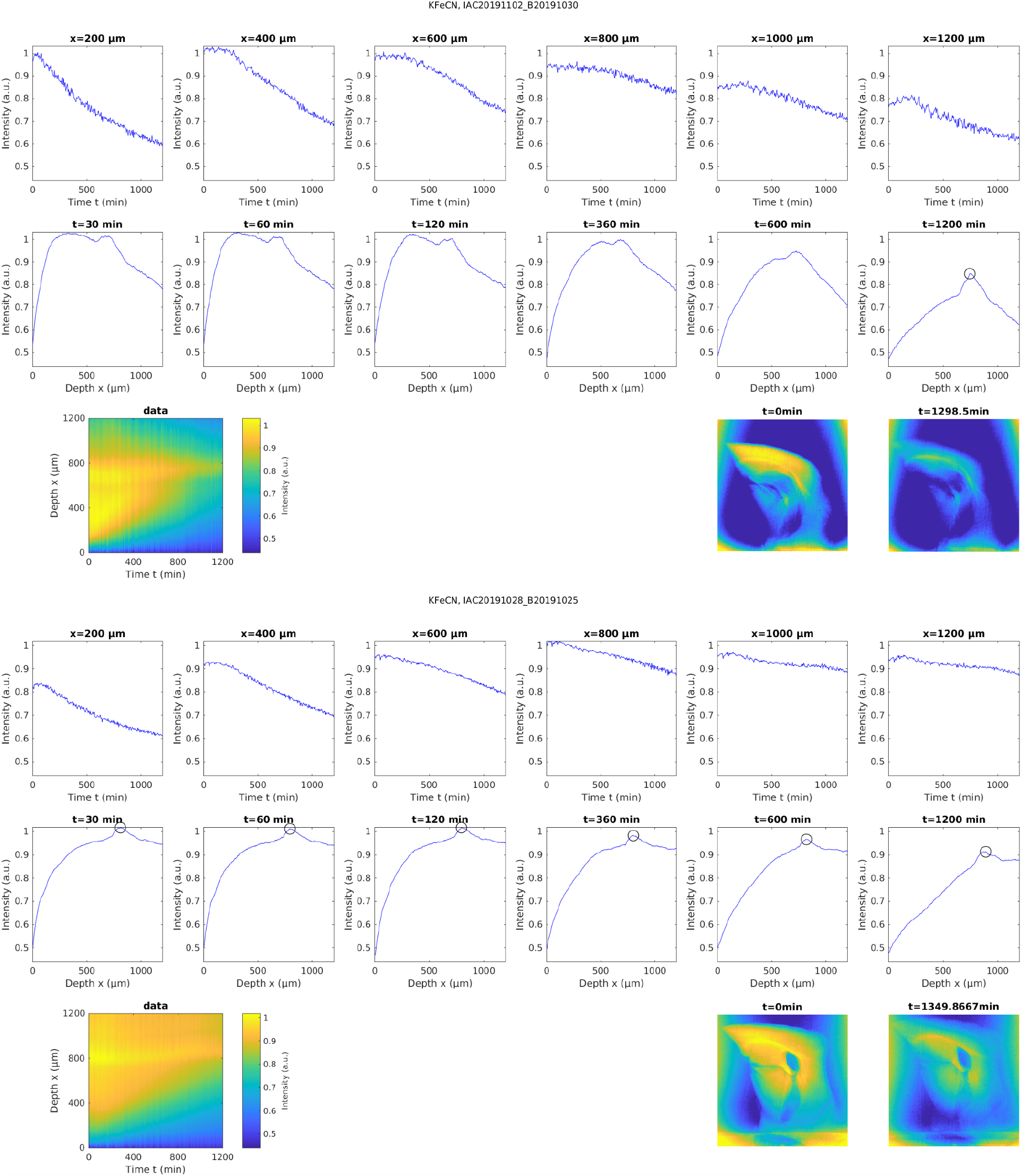
Experimentally measured washout of heavy metals in buffered 2.5% K_4_[Fe(CN)_6_] solution in n=6 different samples that have previously been stained with buffered OsO_4_ for 22 hrs. The first row of every sample shows the temporal profile of osmium washout at different cortical depths, while the second row shows the spatial profile of osmium washout at different times after staining onset. The black circles indicate the location of axon tracts. The third row shows the measured spatial-temporal washout of osmium (left) and the corresponding distribution of osmium in projection images of the sample at the beginning of the experiment and after about 22 hours of incubation.

**Supplementary Figure 6:**
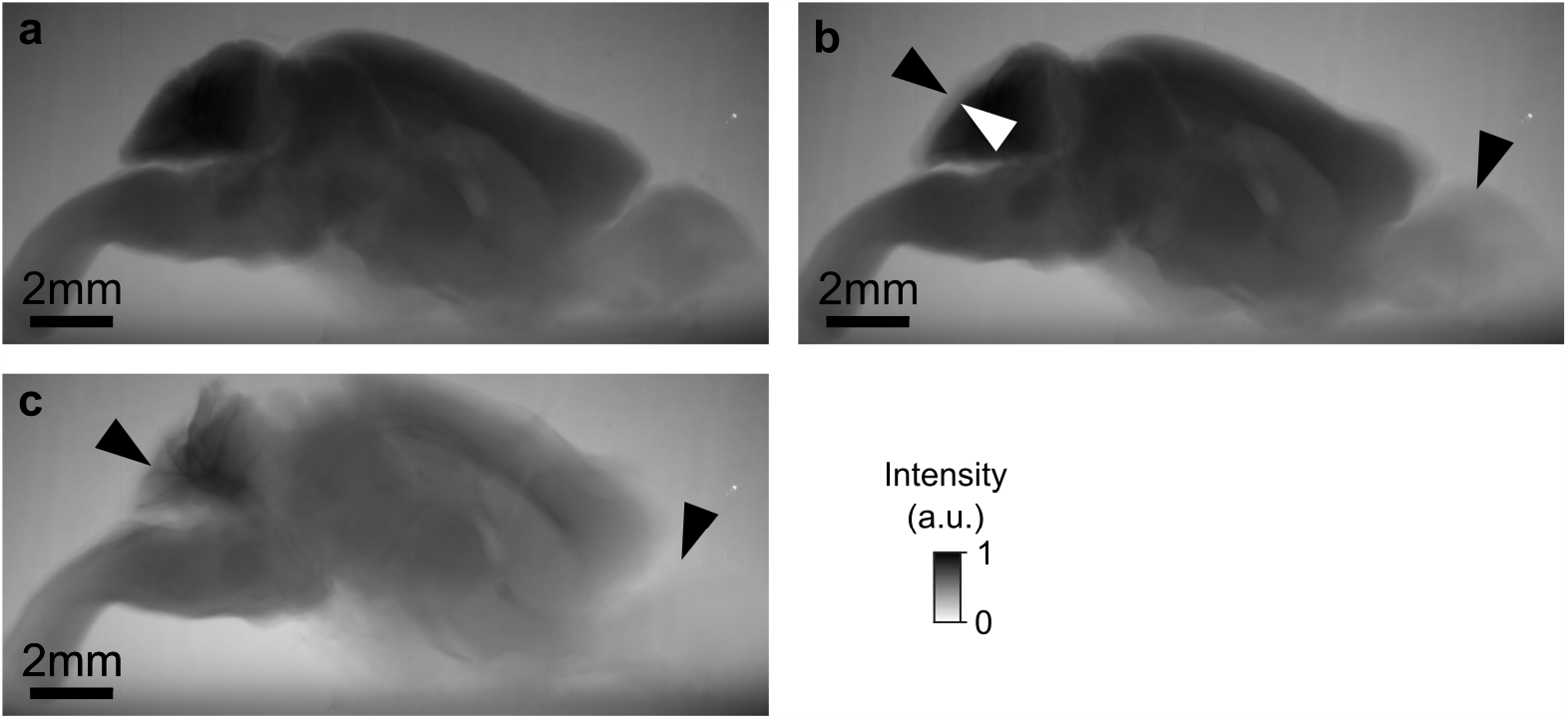
Long incubation in potassium ferrocyanide dissolves and disintegrates the tissue. **a** Whole mouse brain from figure 1d after long incubation in osmium. **b** Subsequently, the osmium solution was replaced by 2.5% buffered potassium ferrocyanide solution and the sample was incubated for 16 hrs. The tissue clearing is particularly well visible in the cerebellum and olfactory bulbs (arrowheads). **c** After another 108 hrs in buffered potassium ferrocyanide a large fraction of the cerebellum and the olfactory bulbs has been dissolved. Scale bar 2 mm.

**Supplementary Figure 7:**
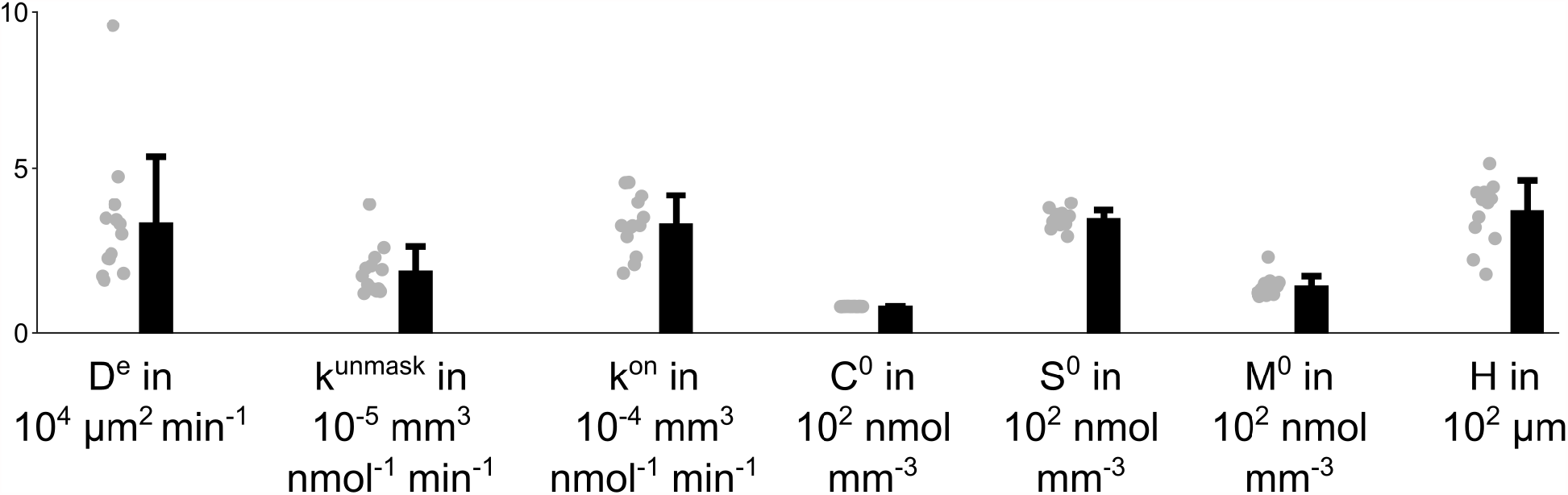
Effective diffusion coefficient D^e^, unmasking rate constant k^unmask^, binding rate constant k^on^, the OsO_4_ concentration in the solution C^0^, the initial density of binding sites S^0^, the initial density of masked binding sites M^0^ and the sample curvature height H for n=13 different 4-mm punches that have been immersed for 22 hrs in 2% buffered OsO_4_ solution. The parameters have been fitted to the experimental data by simulating the model equations (2)-(5) and (7), while C^0^ has been measured for each sample individually.

